# Formation of Chromosomal Domains in Interphase by Loop Extrusion

**DOI:** 10.1101/024620

**Authors:** Geoffrey Fudenberg, Maxim Imakaev, Carolyn Lu, Anton Goloborodko, Nezar Abdennur, Leonid A. Mirny

**Affiliations:** Graduate Program in Biophysics, Harvard University, Cambridge, Massachusetts, USA.; Department of Physics, Massachusetts Institute of Technology (MIT), Cambridge, Massachusetts, USA.; Program for Research in Mathematics, Engineering and Science for High School Students (PRIMES) and Undergraduate Research Opportunities Program (UROP), MIT, Cambridge Massachusetts, USA.; Program in Computational and Systems Biology, MIT, Cambridge, Massachusetts, USA.

## Abstract

Topologically Associating Domains (TADs) are fundamental structural and functional building blocks of human interphase chromosomes, yet mechanisms of TAD formation remain unknown. Here we propose that loop extrusion underlies TAD formation. In this process, cis-acting loop-extruding factors, likely cohesins, form progressively larger loops, but stall at TAD boundaries due to interactions with boundary proteins, including CTCF. Using polymer simulations, we show that this model can produce TADs as determined by our analyses of Hi-C data. Contrary to typical illustrations, each TAD consists of multiple dynamically formed loops, rather than a single static loop. Our model explains diverse experimental observations, including the preferential orientation of CTCF motifs, enrichments of architectural proteins at TAD boundaries, and boundary deletion experiments, and makes specific predictions for depletion of CTCF versus cohesin. The emerging picture is that TADs arise from actively forming, growing, and dissociating loops, presenting a framework for understanding interphase chromosomal organization.

## Introduction

Interphase chromosome organization underlies critical cellular processes, including gene regulation via enhancer-promoter interactions in three dimensions. Recent advances in mapping chromosomal interactions genome-wide have uncovered that interphase chromosomes of higher Eukaryotes are partitioned at the sub-megabase scale into a sequence of self-interacting TADs (Dixon et al., 2012; Nora et al., 2012), or domains (Rao et al., 2014; Sexton et al., 2012). An increasing number of studies have found important functional roles for TADs in the control of gene expression and development (Andrey et al., 2013; Lupiáñez et al., 2015; Symmons et al., 2014).

TADs are contiguous regions of enriched contact frequency that appear as squares in a Hi-C map (**Fig 1A**), and are relatively insulated from neighboring regions. Many TADs have homogeneous interiors, while others have complex and hierarchical structures, and particularly sharp or enriched boundaries (**fig. S1**). More recently, high resolution maps revealed peaks of interactions between loci at the boundaries of TADs (“peak-loci” (Rao et al., 2014)).

**Fig 1.**
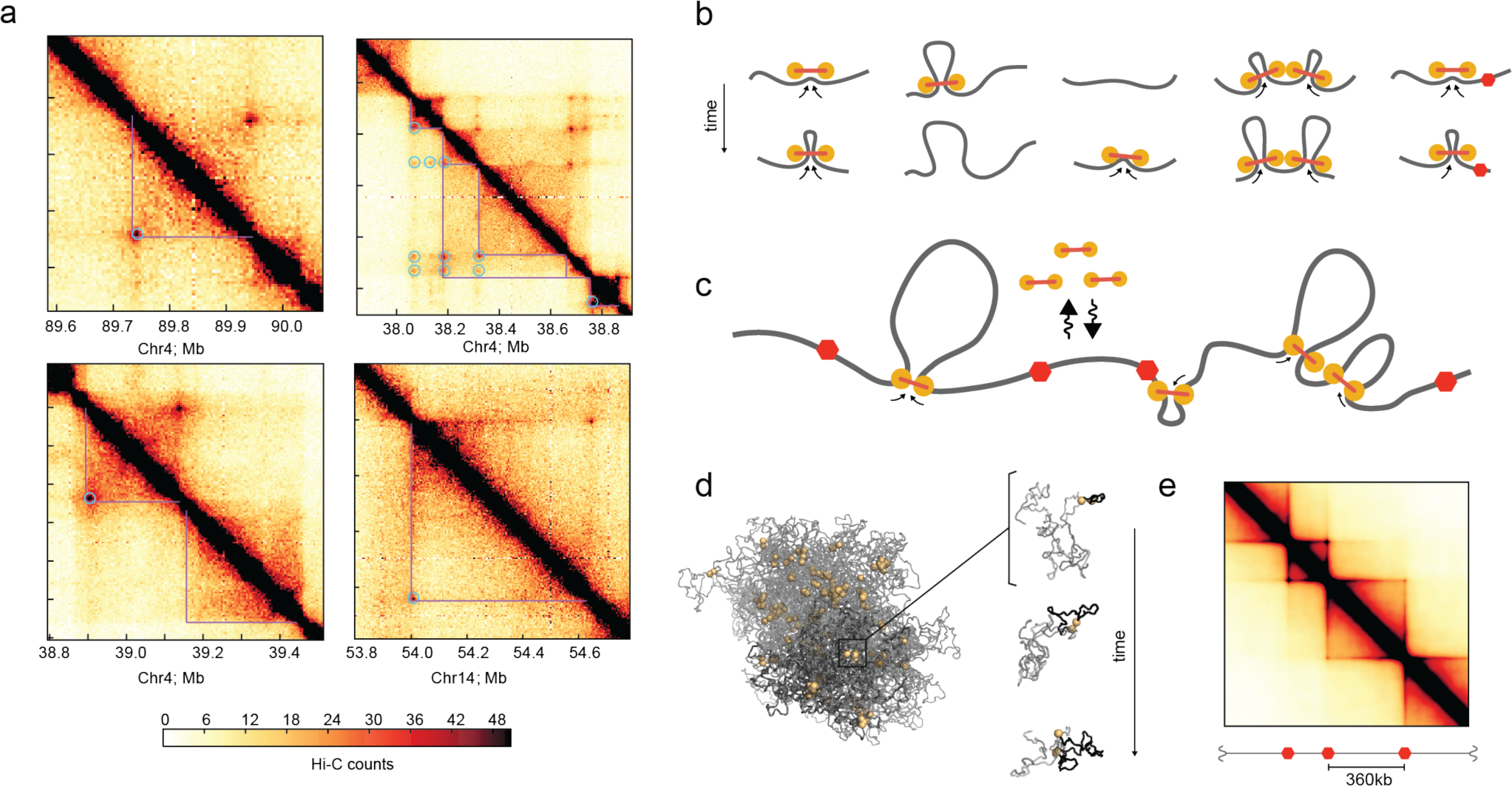
Loop extrusion as a mechanism TAD formation. **a.** Examples of Hi-C contact maps at 5kb resolution showing TADs from four chromosomal regions (GM12878 in-situ MboI (Rao et al., 2014)), highlighting TADs (purple lines) and interaction peaks (blue circles). **b.** Model of LEF dynamics, LEFs shown as linked pairs of yellow circles, chromatin fiber in grey. From left to right: extrusion, dissociation, association, stalling upon encountering a neighboring LEF, stalling at a BE (red hexagon). Also see **fig. S2.** **c.** Schematic of LEF dynamics (**Movie-M1, Movie-M2**). **d.** Conformations of a polymer subject to LEF dynamics, with processivity 120kb, separation 120kb. *Left*: shows LEFs (yellow), and chromatin (grey), for one conformation, where darker grey highlights the combined extent of three regions of sizes (180kb, 360kb, 720kb) separated by BEs. *Right*: shows the progressive extrusion of a loop (black) within a 180kb region. **e.** Simulated contact map for processivity 120kb, separation 120kb.

TADs differ from larger-scale compartments in many regards. First, they do not necessarily form an alternating ‘checkerboard’ pattern of enriched contact frequencies (Lajoie et al., 2014). Often, several TADs reside within a single contiguous compartment (Gibcus and Dekker, 2013; Gorkin et al., 2014). TADs are largely stable across cell-types, while compartments are not (Dekker and Heard, 2015). TADs and compartments display distinct responses to biological perturbations, including cohesin knockdown (Seitan et al., 2013), and distinct relationships with other nuclear landmarks, including the lamina (Kind et al., 2015). Although correlated with replication timing, available evidence indicates that TADs are independent of DNA replication itself, they: appear early in G1, before replication has a chance to influence chromosome organization (Dileep et al., 2015; Naumova et al., 2013); are maintained in cells were replication has been halted by thymidine block (Naumova et al., 2013); and remain present, though are on-average weaker, in senescent cells that do not replicate their DNA (Chandra et al., 2015).

Although often illustrated as large loops, several lines of evidence indicate that TADs are not simply stable loops formed between two boundary loci. First, only 50% of TADs have corner-peaks (Rao et al., 2014). Second, boundary loci do not appear to be in permanent contact either by FISH (Rao et al., 2014), or by their relative contact frequency (see results). Third, while TADs are enriched in contact probability throughout the domain, polymer simulations show that simple loops display enrichment only at the loop bases, unless the loop is very short (Benedetti et al., 2014; Doyle et al., 2014). For these reasons, identifying mechanisms of how TADs are formed, and what constitutes a TAD boundary, remain important open questions.

While polymer models have provided insight into multiple levels of chromosome organization (Bau et al., 2011; Lieberman-Aiden et al., 2009; Marko and Siggia, 1997; Naumova et al., 2013; Rosa and Everaers, 2008), relatively few have focused on TADs. Of those that have considered TADs, some have focused primarily on characterizing chromosome structure rather than mechanisms of folding (Giorgetti et al., 2014; Hofmann and Heermann, 2015). Others (Barbieri et al., 2012; Jost et al., 2014) have considered models where monomers of same type experience preferential pairwise attractions to produce TADs; such models, however, when generalized to the genome-wide scale would require a separate factor to recognize and compact each TAD. With only several types of monomers, this would produce checkerboard patterns for each type, which is characteristic of compartments rather than TADs. One proposed mechanism giving good agreement to the observed TAD organization relies on supercoiling (Benedetti et al., 2014), though the role of supercoiling in eukaryotes remains unclear, and it is not clear if this mechanism can also give rise to corner-peaks between loci at the boundaries of TADs.

Here we propose a mechanism whereby TADs are formed by loop extrusion (Alipour and Marko, 2012; Nasmyth, 2001). In this process, cis-acting loop-extruding factors (LEFs, likely cohesins) form progressively larger loops, but are stalled by boundary elements (BEs, including bound CTCF) at TAD boundaries (**Fig. 1B-C**). We tested this mechanism using polymer simulations of the chromatin fiber subject to the activity of LEFs. We find that loop extrusion can indeed produce TADs that qualitatively and quantitatively agree with our new analyses of the highest-resolution Hi-C data. Moreover, we find that TADs in agreement with Hi-C data are formed by the dynamics of forming, growing and dissociating loops. Importantly, our work provides a mechanism for preferentially forming loops within TADs, and not between TADs; such a mechanism is implicitly assumed in structural models of TADs formed by dynamic loops (Giorgetti et al., 2014; Hofmann and Heermann, 2015). Loop extrusion (Alipour and Marko, 2012), first introduced as processive loop enlargement by condensin (Nasmyth, 2001), has been implicated in mitotic chromosome compaction (Goloborodko et al., 2015; Naumova et al., 2013), and chromosome segregation in bacteria (Gruber, 2014; Wang et al., 2015). Still, these previous proposals did not consider any role of loop extrusion for TAD formation in interphase and did not directly test the impact of loop extrusion on 3D spatial organization. We note that a similar hypothesis has been put forward but not tested by simulations (Nichols and Corces, 2015), and a study investigating a very similar loop extrusion model was published while our manuscript was available as a preprint (http://dx.doi.org/10.1101/024620) and under review (Sanborn et al., 2015).

## Results

### Model

To demonstrate how the mechanism of loop extrusion can lead to the formation of TADs, we modeled the process by coupling the 1D dynamics of loop extrusion by LEFs with polymer dynamics in 3D (**Fig. 1B-C, fig. S2**). Upon binding to the chromatin fiber, a LEF holds together two directly adjacent regions; it then extrudes a loop by translocating along the chromatin fiber in both directions, holding together progressively distant regions of a chromosome (**Movie-M1, Movie-M2**). Translocation stops when the LEF encounters an obstacle, either another LEF, or a BE; if halted only on one side, LEFs continue to extrude on the other side. Throughout this process, LEFs can stochastically dissociate, releasing the extruded loop; for generality, we assume that this occurs uniformly across the genome.

Boundary elements (BEs) are the second crucial component underlying the formation of TADs in our proposed mechanism. BEs are fixed genomic loci that stall LEF translocation and ensure that extruded loops do not cross TAD boundaries. BEs in our model correspond to any impediment to loop extrusion. As such, BEs *in vivo* might be formed by specifically bound architectural proteins (e.g. CTCF), or a generally high occupancy of architectural proteins (Van Bortle et al., 2014). Active promoters bound by bulky transcription-associated machinery may also form BEs; indeed, promoters are known to be enriched at TAD boundaries (Dixon et al., 2012). The combined action of LEFs and BEs leads to enrichment of interactions within TADs and effective insulation between neighboring TADs. Note that the interactions induced by LEFs cannot simply be described by an effective pairwise potential, as their action is time-dependent and multi-point in nature (see **Methods**).

To consider how loop-extrusion dynamics can spatially organize a chromosome in 3D, we model a 10Mb region of the chromatin fiber as a polymer subject to the activity of associating and dissociating LEFs (**Fig 1C**). We model the chromatin fiber as a series of 10nm monomers, each representing roughly three nucleosomes, ∼600bp (Naumova et al., 2013). As previously (Naumova et al., 2013), the polymer has excluded volume interactions, has no topological constraints and is simulated by Langevin dynamics using OpenMM (Eastman et al., 2013). LEFs impose a system of bonds on the polymer: a bound LEF forms a bond between the two ends of an extruded loop, and the bond is re-assigned to increasingly separated pairs of monomers as a LEF translocates along the chromosome; when a LEF unbinds, this bond is removed. BEs, which halt LEFs translocation, were placed at fixed positions with sequential separations of 180kb, 360kb, and 720kb, through the 10Mb region.

The dynamics of loop extrusion are determined by two independent parameters (**Fig 2B, fig. S2**): the average linear *separation* between bound LEFs, and the LEF *processivity,* i.e. the average size of a loop extruded by an unobstructed LEF over its lifetime (Goloborodko et al., 2015). Our model is additionally characterized by parameters governing the diffusivity of chromatin, polymer stiffness, density, and the Hi-C capture radius (**Methods**). For each set of parameter values, we ran polymer simulations long enough to allow many association/dissociation events per LEF (10-160 events, **Movie-M1, Movie-M2**). From simulations, we obtain an ensemble of chromosome conformations (**Fig 1D**) and compute contact frequency maps (“simulated Hi-C”, **Fig 1E**) that can be compared with experimental Hi-C data.

**Fig 2.**
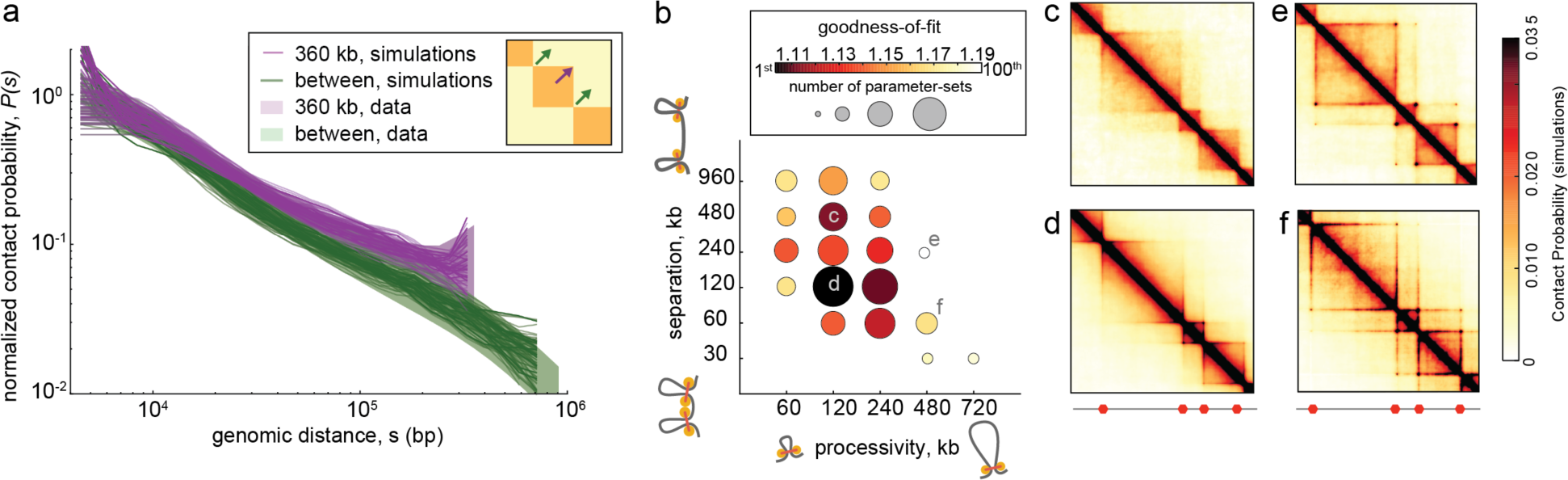
Quantitative analysis of loop extrusion. **a.** Experimental *P(s)* (shaded areas) versus simulated *P(s)* for the 100 best-fitting parameter-sets (lines, one per parameter-set) within TADs (purple) and between TADs (green). Experimental *P(s)* calculated from 2kb contact maps and normalized to one at 4kb; shaded area shows 10^th^ and 90^th^ percentiles at each genomic distance. Simulated *P(s)* shown with vertical offsets obtained from fitting procedure (**Methods**). **b.** Goodness-of-fit versus LEF processivity and separation for the 100 best-fitting parameter-sets (from 6912 total parameters-sets). Circle areas represent the number of parameter-sets among the top-100, while color quantifies the best-fit at each *processivity-separation* pair; a value of 1 indicates a perfect fit. **c-f.** Simulated contact maps for the indicated *processivity-separation* pairs.

### TADs from Loop Extrusion

For a range of LEF processivities and separations, we observe the formation of TADs on a simulated Hi-C map (**Fig 2C-E**). In our model, TADs emerge due to the following process: Due to the stalling of LEF at BEs, loops of various sizes dynamically form in the region between neighboring BEs (**fig. S3**). In turn, direct contacts between loop-bases preferentially enrich the contact frequencies of different sets of loop-bases in different cells in these regions. Averaged over a large population of cells, this mechanism results in TADs as observed in Hi-C data. For some parameter values we observe formation of homogenous TADs; other simulated parameter sets lead to the formation of peaks at corners of TADs, or enrichment of contacts at the boundary of TADs (**Database D1**). These diverse architectural features are similar to those observed in experimental data (**fig. S1**).

In addition to agreement with architectural features, our model naturally recapitulates the results of TAD boundary deletion experiments (Nora et al., 2012). In experiments, upon deletion of a TAD boundary, the TAD spreads to the next boundary; this indicates that preferential interactions between loci in a TAD are not hard-wired, and that boundary elements play crucial roles. Consistently, in our model, deletion of a BE leads to spreading of a TAD until the next BE (**fig. S4**).

We next tested the ability of our model to reproduce an important quantitative characteristic of TAD organization observed in Hi-C data: the dependence of Hi-C contact frequency *P(s)* with distance *s*, used previously for quantifying polymer models (Barbieri et al., 2012; Benedetti et al., 2014; Le et al., 2013; Naumova et al., 2013; Rosa et al., 2010). We aim to reproduce both *P(s)* within TADs (separately for TAD sizes 180kb, 360kb and 720kb) and *P(s)*between TADs, which is ∼1.5-fold smaller and scales differently with distance (**Fig 2A, fig. S5**). For each of 6912 parameter-sets, we determined the goodness-of-fit as the geometric standard deviation between the four experimental and simulated *P(s)* curves (**Methods**). For each pair of values of LEF processivity and LEF separation, we quantified the best achieved goodness-of-fit and the number of times a pair appears among the top-100 parameter-sets (**Fig 2B**).

We found that the best agreement with Hi-C data is achieved for LEF processivity of ∼120-240kb and LEF separation ∼120Kb (**Fig 2A-B**), where the resulting TADs consist of dynamically forming, growing and dissociating loops (**Fig 3A, fig. S3**). In this regime, LEFs extrude ∼120Kb loops relatively independently, as there are substantial gaps between LEFs (30-72% coverage of TADs by loops). Notably, despite a ∼2-fold depletion of contact probability between neighboring TADs, polymer conformations display high spatial overlap between adjacent TADs rather than appearing as segregated globules in this regime (**Fig 3B, fig. S6C**).

**Fig 3.**
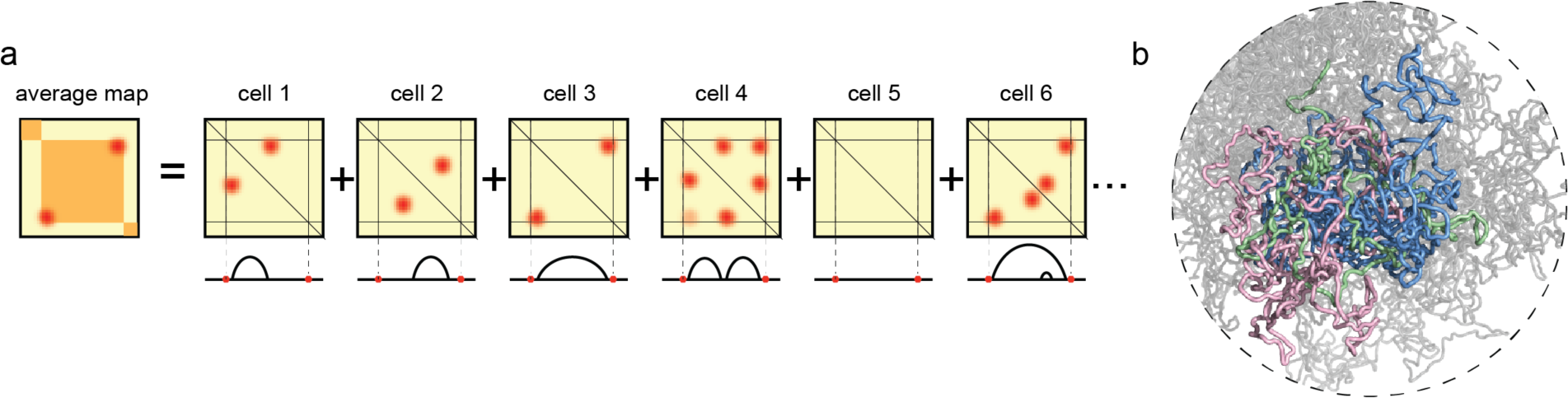
TADs formed by LEFS consist of dynamically forming, growing and dissociating loops. **a.** Illustration of how TADs formed by loop extrusion result from averaging the dynamic positions of loop-bases over many cells, including configurations with nested (cell 6) and consecutive (cell 4) loops (also see **fig. S3**). **b.** Conformation of a polymer subject to LEF dynamics with processivity 120kb, and separation 120kb. Three neighboring regions between BEs of sizes (180kb, 360kb, 720kb) colored in (green, pink, blue). Contacts from an ensemble of such conformations are averaged together to form a contact map.

Our simulations make predictions about changes in contact maps and spatial distances that would result from experimental perturbations of architectural proteins (**fig. S7**). For example, we predict that reducing the strength of BEs (e.g. by CTCF depletion) would reduce insulation between neighboring TADs yet would have little effect on spatial distances of loci within TADs and only moderately reduce spatial distances of loci between TADs. Increasing the separation of LEFs (e.g. by cohesin depletion) would also make TADs weaker. In contrast, however, this perturbation would be accompanied by more drastic increases in spatial distances of loci both within and between TADs.

### TAD Corner Peaks

An important feature of Hi-C TADs is that many appear to have peaks of interactions at their corners (∼50%, (Rao et al., 2014)). Our analysis of Hi-C data shows that TADs with and without peaks have similar *P(s)*, suggesting a similar underlying organizational mechanism, independent of the corner-peak (**fig. S5**). In agreement, our model shows that the mechanism of loop extrusion can produce both types of TADs, as increasing LEF processivity naturally strengthens peaks at TAD corners while LEF dynamics still provide within-TAD enrichment (**Fig 2E-F, fig. S8**). Interestingly, our simulations show that TADs with visibly strong peaks do not require permanent contact between BEs, in agreement with our analyses of Hi-C data (**fig. S8**).

Indeed, we find that strong loops between BEs provide among the worse fits to Hi-C data, with exceedingly-strong corner-peaks and a lack of visible TADs (**Fig 4, fig. S9B**). Our results indicate that that a single stable loop does not describe TADs as observed in Hi-C, building on previous results (Benedetti et al., 2014; Doyle et al., 2014; Hofmann and Heermann, 2015), and standing in contrast with popular depictions of TADs as loops (Rao et al., 2014). Instead, our model predicts that TADs result from the activity of LEFs in the region between BEs, whereas corner-peaks emerge when LEFs transiently form BE-to-BE loops; these predictions can be tested with high-resolution live-cell imaging.

**Fig 4.**
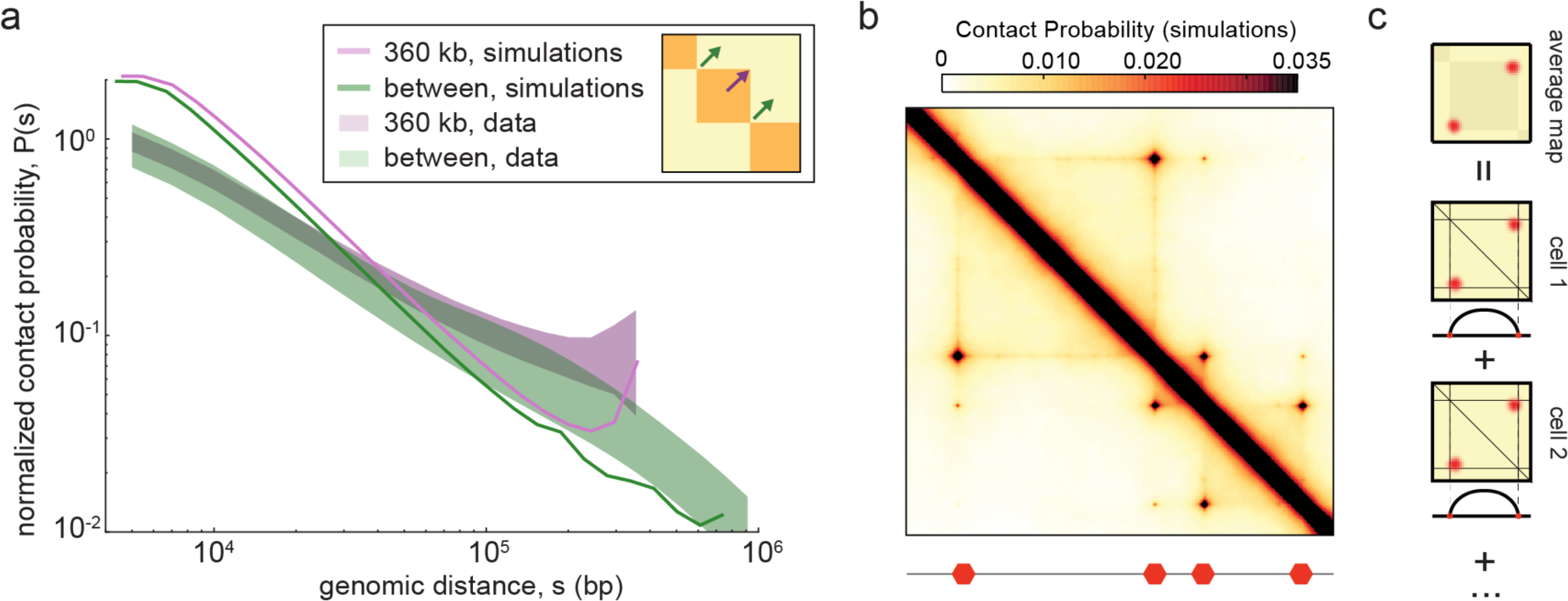
Simple Strong Loops are not TADs. **a.** Experimental *P(s)* (shaded areas) versus simulated *P(s)* (solid lines) for a parameter-set with a strong loop between neighboring BEs, calculated as in **Fig 2A**; in this case, the fit is relatively poor (1.4137, rank 2208 out of 6912). We note that with these parameters loops are not completely permanent and BEs are in contact 27% of the time for the smallest 180kb TAD, and 14% for the largest 720kb TAD. **b.** Simulated contact map for a simple strong loop with processivity 960kb, separation 960kb. **c.** Illustration of how a single loop present in many cells leads to strong corner-peaks between neighboring BEs.

### TADs require long-range insulation

Importantly, insulation between neighboring TADs in our model does not arise from direct physical blocking of interactions between distal genomic regions by BEs. Instead, our model relies on the ability of BEs to regulate the translocation of LEFs. LEFs allow for insulation to be mediated over spatial and genomic distances much larger than the physical size of the BE. To rule out the possibility that a bulky BE is sufficient to insulate neighboring TADs, we performed simulations of this scenario. Indeed, in simulations where a BE is simply a bulky object, we see no long-range insulation and fail to obtain TADs (**fig. S10**). Similarly, in simulations where the chromatin fiber is locally very stiff at a BE, we again only see local insulation, and also fail to obtain TADs (**fig. S10**). Together, these simulations highlight the role of LEFs for imposing insulation at the scale of whole TADs.

Another important characteristic of our model is that loops extruded by LEFs act in cis, along the chromatin fiber, and do not impose interactions between genomically distal loci or loci on different chromosomes. Indeed, when we analyzed the interaction patterns of peak-loci in Hi-C data, we found that there was no enrichment of contacts between pairs of peak-loci at larger separations on the same chromosome or between different chromosomes (**fig. S11**). This pattern is consistent with our model, but is inconsistent with models that rely on direct interactions between BEs when such loci come into spatial proximity.

To rule out the mechanism whereby TADs are formed by direct BE-to-BE associations, we performed simulations where any two BEs would interact when they came into close spatial proximity (**fig S11**). Biologically, this represents a scenario where proteins interact to bridge cognate genomic elements (Barbieri et al., 2012; Bohn and Heermann, 2010; Brackley et al., 2015; Scolari and Lagomarsino, 2015), for example via interactions mediated by dimerization of bound CTCF. Our simulations confirmed that a direct BE-to-BE mechanism has no way of distinguishing between distant or proximal chromosomal regions; instead, all pairs of BEs display peaks of contact probability. Moreover, direct BE-to-BE interactions alone imposed negligible insulation between neighboring TADs, even in the case of strongly interacting BEs. Together, these results demonstrate the utility of LEFs stalled by BEs for restricting potentially interacting pairs of loci to those that are within TADs.

### Complex TAD architectures

LEF dynamics *in vivo* can have additional subtleties beyond those in the minimal model introduced above. For example, the lifetimes of LEFs stalled at BEs may be different from moving LEFs; LEFs may backtrack or pause; there may be several types of LEFs with different processivities; and LEFs may be actively loaded or unloaded at particular elements. To understand how locus-specific parameters of LEF dynamics can form a basis for the rich variety of TAD domain architectures *in vivo*, we considered three particular extensions of our minimal model.

First, we investigated simulations with semi-permeable boundary elements, which reflects either BEs that are present in a fraction of cells, or BEs that probabilistically stall LEF translocation. We found this can reproduce a nested TAD-in-TAD organization (**Fig 5A**). Second, we examined simulations where LEFs are preferentially loaded at one side of a TAD, as might result from locus-specific positioning of loading factors. We found this could reproduce an asymmetric TADs with stronger interactions between the BE and the body of the TAD (**Fig 5B**).

**Fig 5.**
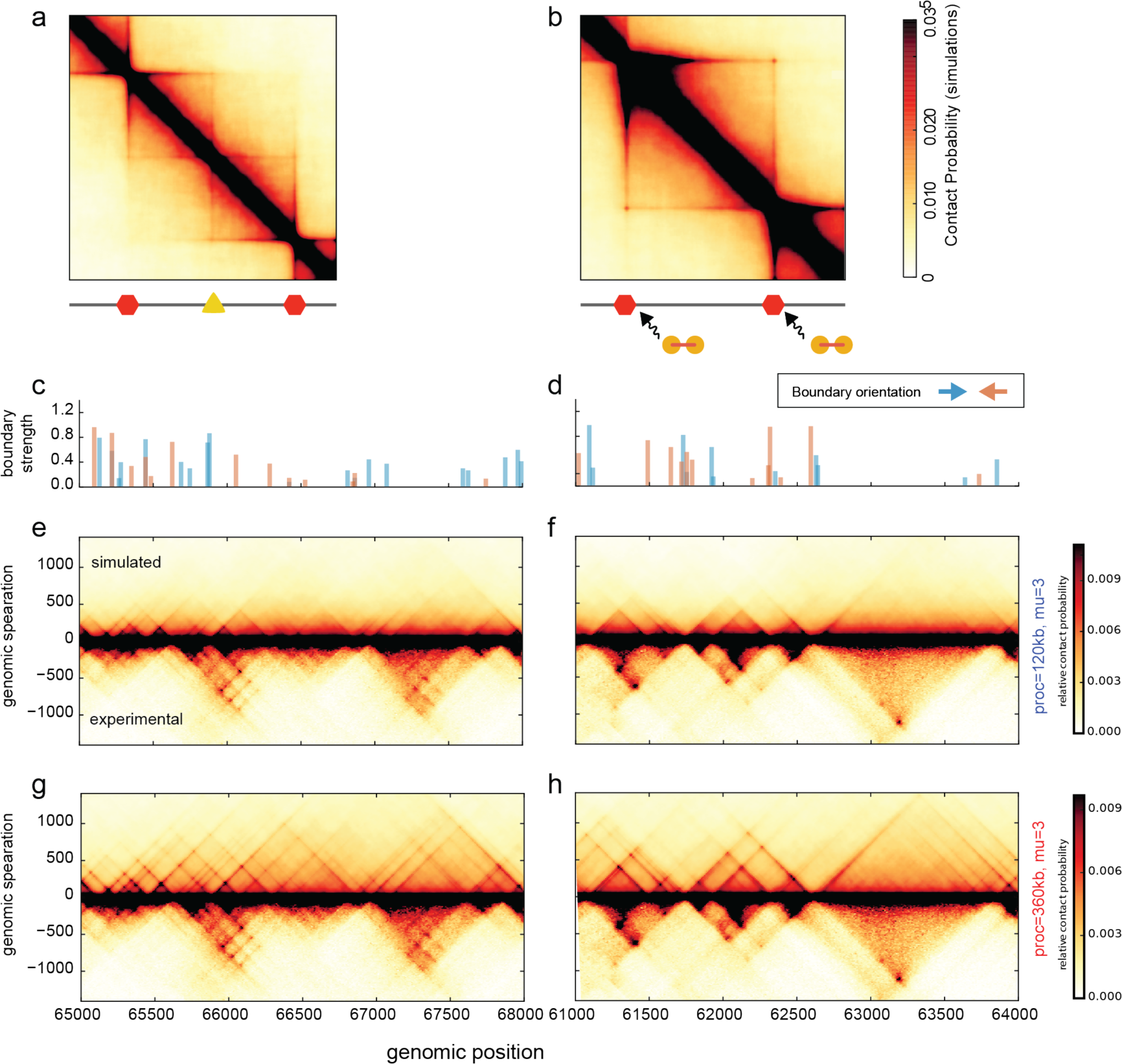
Complex TAD architectures from loop extrusion. **a.** Simulated contact map for a model with a semi-permeable BE (yellow triangle), displaying nested TADs. **b.** Simulated contact map for a model with asymmetric loading of LEFs. **c-d.** Orientation-specific BE strength profile for regions simulated in e-h; 12kb resolution. BE strength reflects the average number of BEs bound within a 12kb (20 monomer) region. **e-h.** Simulated contact maps for regions of human chr14, GM12878 cell type, for models with orientation specific BEs of varying permeability. Maps are compared with experimental maps for the same regions at the same 12kb resolution (also see **Methods, fig. S12**). LEF processivity is 120kb (e,f) and 360kb (g,h).

Finally, we investigated a complex system of orientation-specific BEs with locus-specific strengths (**Fig 5C-H**). For these simulations, we converted ChIP-seq data for CTCF and cohesin over a 15Mb region of human chromosome 14 into BE occupancy and directionality (**Methods**). We found that when parameters of the minimal model described above were used, the abundance of semi-permeable boundaries in this system produced contact maps with good agreement to Hi-C maps at short distances (<400kb), but poor agreement at further distances (**fig. S12**). We found that better agreement at far distances can be obtained if we increased LEF processivity to 360kb (corresponding to an increase in average loop size from ∼75kb to ∼135kb).

Still, even with increased LEF processivity, agreement along the chromosome was non-uniform (**fig. S12**). Surprisingly, this non-uniform agreement was not seen in (Sanborn et al., 2015). Our observation of non-uniform agreement along the chromosome possibly reflects: additional undetermined factors underlying BEs, locus-specific details of LEF dynamics (including loading and unloading); the role of higher-order active and inactive compartments (Brackley et al., 2015; Jost et al., 2014); or locus-specific experimental details of Hi-C and ChIP-seq (Imakaev et al., 2012; Yaffe and Tanay, 2011). Nevertheless, our simulations show that loop extrusion with BEs imposed by bound CTCF constitutes a useful step towards understanding the fine structure of *in vivo* contact maps.

## Discussion

In the context of loop extrusion as a mechanism underlying TAD formation, certain architectural proteins are attractive molecular candidates for LEFs and BEs; establishing these roles would both explain existing experimental results, as well as suggest new experiments. Generally, Structural Maintenance of Chromosome (SMC) complexes, including cohesin and condensin, with a hypothesized motor function and a similar molecular architecture to known motor proteins (Guacci et al., 1993; Nasmyth, 2001; Peterson, 1994), are plausible candidates for LEFs. Additionally, both cohesin and condensin complexes have been hypothesized to have the ability to extrude chromatin loops (Alipour and Marko, 2012; Nasmyth, 2001). In interphase, cohesins have been implicated in TAD organization (Mizuguchi et al., 2014; Sofueva et al., 2013; Zuin et al., 2014) and chromatin loops (Kagey et al., 2010) beyond their role in sister chromatid cohesion, and have been observed to dynamically bind to chromatin, even before replication (Gerlich et al., 2006). Additionally, cohesin is enriched at interphase TAD boundaries (Dixon et al., 2012) and loops (Rao et al., 2014), and its depletion makes TADs less prominent (Sofueva et al., 2013; Zuin et al., 2014). Finally, increasing cohesin binding time by depleting the cohesin unloader Wapl (Tedeschi et al., 2013) condenses interphase chromosomes into a prophase-like ‘vermicelli’ state; our model predicts this behavior if LEF processivity is greatly increased (**fig. S13**).

Boundary-elements in our model correspond to any impediment to loop extrusion; CTCF is a particularly relevant molecular candidate (Hou et al., 2008; Ong and Corces, 2014). CTCF is enriched at TAD boundaries (Dixon et al., 2012; Rao et al., 2014; Rudan et al., 2015; Zuin et al., 2014), its depletion makes TADs less prominent (Zuin et al., 2014), and it has a relatively long residence time on chromatin (Nakahashi et al., 2013). In addition, analogous to LEF accumulation at BEs in our model (**Fig 6B**), we predict that cohesin accumulates at CTCF binding sites, but only when CTCF is bound at these sites (Parelho et al., 2008).

**Fig 6.**
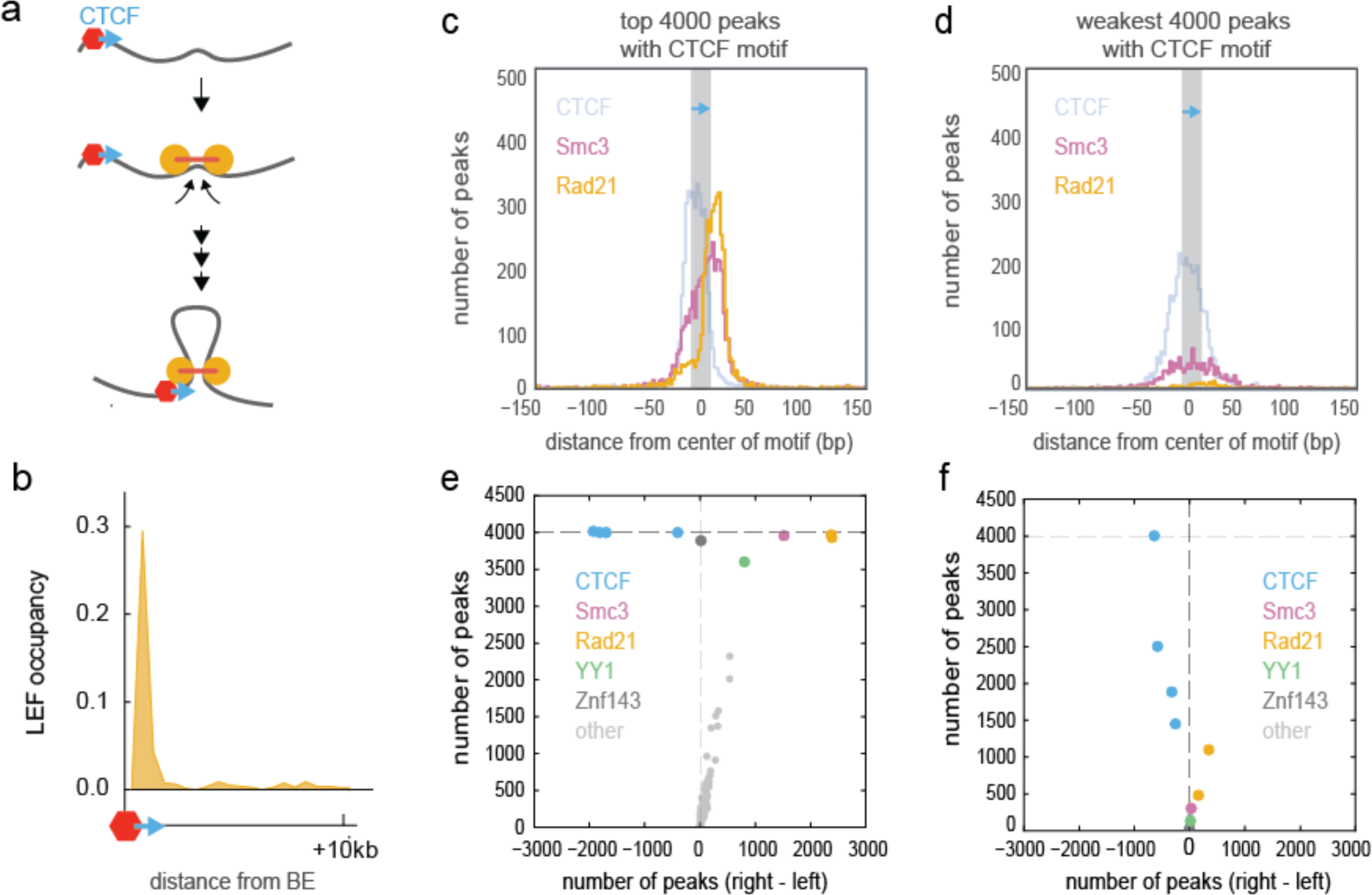
CTCF as an orientation-dependent boundary element. **a.** Loop extrusion and an orientation-dependent boundary function of bound CTCF elements can lead to enrichment of inward-oriented CTCF sites at TAD boundaries, even over large genomic distances (see **fig. S14**). **b.** Accumulation of LEFs at BEs for simulations with processivity 120kb and separation 120kb. **c.** Distributions of CTCF, Smc3, and Rad21 ChIP-seq peak summits in the vicinity of the 4000 strongest CTCF binding peaks with a detected CTCF motif instance in GM12878. The peak summits are oriented relative to the center of the nearest CTCF motif instance. All motif instances are oriented in the same direction, depicted by the blue arrow. **d.** Same, but for the weakest 4000 motif-associated CTCF binding sites. **e.** Asymmetry in ChIP-seq peaks of ENCODE factors around the strongest 4000 CTCF ChIP peaks with a detected CTCF motif instance. Each dot represents a GM12878 ChIP-seq track. The y-axis shows the number of peaks of a factor found within +/- 200 bp of a CTCF motif instance. The x-axis shows the difference between the number of factors found on the right of the center of the motif and on the left, i.e. asymmetry of the factor relative to a CTCF motif. CTCF in blue, Smc3 in magenta, Rad21 in orange, ZNF143 in dark grey, YY1 in green, other factors in light grey. **f.** Same, but for the weakest 4000 motif-associated CTCF ChIP peaks.

Finally, if CTCFs halt loop extrusion and stabilize loops in an orientation-dependent manner, then the mechanism of loop extrusion predicts the observed enrichment in convergent CTCF sites at TAD boundaries and loop bases (Rao et al., 2014; Rudan et al., 2015) even at very large genomic separations (**Fig 6A, fig. S14**), as well as the results of manipulating CTCF site orientation (Guo et al., 2015; Narendra et al., 2015; Sanborn et al., 2015; de Wit et al., 2015). Indeed, CTCF binding sites are oriented such that the C-terminus of bound CTCF (Nakahashi et al., 2013), known to interact with cohesin (Xiao et al., 2011), faces the interior of TADs. The interaction of CTCF and cohesin may stabilize cohesin-mediated loops either directly or by shielding cohesins from the unloading action of SA2-interacting Wapl (Ouyang et al., 2013), similar to the interactions of shugoshin (Hara et al., 2014) and sororin (Nishiyama et al., 2010) with cohesin. Consistently, we find that cohesin ChIP-seq peaks are enriched in this exact orientation-dependent manner around strongly enriched CTCF peaks (**Fig 6C-F**). Taken together, these observations support a mechanism where CTCF acts as a boundary element that impedes loop extrusion in an orientation-dependent manner.

With these molecular roles, we predict that depletion of either CTCF or cohesin from chromosomes would disrupt TADs, but would differentially affect spatial distances (**fig. S7**). Depletion of CTCF would not affect distances between loci within TADs, but would decrease distances between neighboring TADs to the within-TAD level. In contrast, depletion of cohesin would increase distances for loci within TADs. Available imaging data supports decompaction following cohesin depletion (Nolen et al., 2013; Sofueva et al., 2013; Zuin et al., 2014), and lack of decompaction following CTCF depletion (Nolen et al., 2013). In contrast, models where cohesin is strictly loaded by CTCF (Nichols and Corces, 2015) predict an equivalent result for either CTCF or cohesin knockdown. However, further validation of our model would require new methods for architectural protein removal, as available techniques have yet to fully disrupt TAD formation. Similarly, new technologies will be required to directly test the mechanism of loop extrusion. One possibility would be to specifically label and track boundary loci using live-cell super-resolution imaging and test whether they display periods of directed motion towards each other. Finally, *in vitro* single-molecule techniques can be used to characterize the subtleties of LEF dynamics, including whether: lifetimes of LEFs stalled at BEs are different from moving LEFs; different classes of LEFs have different processivities; and LEFs may be actively loaded and unloaded at particular elements.

The mechanism of loop extrusion in interphase has additional, potentially far-ranging, consequences for processes in the nucleus. First, enhancer-promoter pairings can be dictated by the relative placement of BEs, including CTCF (Hou et al., 2008). Second, loop extrusion may have an even stronger effect if LEFs stall at promoters, effectively turning the enhancer-promoter search process into a 1D search process. Third, loop extrusion may facilitate high-fidelity VDJ and class switch recombination and other processes dependent on long-range intra-chromosomal looping with specific orientations, particularly given the observed interplay between CTCF and cohesin (Alt et al., 2013; Degner et al., 2011; Dong et al., 2015; Lin et al., 2015).

Finally, the mechanism of TAD formation via loop extrusion studied here is similar to the proposed mechanism of mitotic chromosome condensation (Alipour and Marko, 2012; Goloborodko et al., 2015, 2016; Nasmyth, 2001; Naumova et al., 2013), but with the addition of BEs and many fewer, less processive, LEFs. Accordingly, increasing the number and processivity of LEFs and removing BEs could underlie the transition from interphase to mitotic chromosome organization. Conversely, upon exit from mitosis, interphase 3D chromosome organization can be re-established simply by restoring BE positions, which could potentially be epigenetically inherited bookmarks (Kadauke and Blobel, 2013).

## Experimental Procedures

### Model Overview

To simulate how the mechanism of loop extrusion can actively compact TADs, we modeled the process by coupling the 1D dynamics of loop extrusion by LEFs with 3D polymer dynamics. We use a discrete model for 1D LEF dynamics, which imposes a system of bonds on the simulated 3D polymer dynamics. This allows us to generate average simulated contact maps for different parameter values in our model, which can in turn be compared with experimental Hi-C contact maps.

### 1D LEF dynamics

LEF translocation along a chromatin fiber was simulated on a 1D lattice. Each lattice position later corresponds to one monomer in the 3D polymer simulation. Each lattice position was characterized by the following parameters: 1. association (birth) probability, 2. dissociation (death) probability.

Since the molecular mechanism of loop extrusion is not known, we model a LEF very generally as having two “heads” connected by a linker, analogously to SMC protein complexes. Each head of a LEF occupied one lattice position at a time, and no two heads could occupy the same lattice position except at birth events.

Birth events occur when the model is initialized, and following each dissociation event; this keeps the total number of bound LEFs constant. In a birth event, a LEF was placed at a random lattice site with probability proportional to the birth probability. With probability 50%, two heads of a LEF were placed to the same location. With probability 50%, we attempted to place the right head of a LEF on the next position (one to the right from the left head), and did so if that site was unoccupied. This possibility was necessary because otherwise LEFs would produce a checkerboard pattern on the contact map at the monomer-scale, as the two heads of a LEF that has never been stalled would always occupy either (even, odd), or (odd, even) positions.

At each time-step, each LEF head translocates to the neighboring lattice site, if that site is not occupied or is not a boundary element (BE). If one head cannot translocate, the other head is unaffected.

At each time-step, a LEF dissociates with a probability equal to the maximum of the death probabilities at the positions of the two heads of a LEF.

In principle, this formulation allows for a very complex description of LEF dynamics where both rates are variable throughout the genome. However, since it remains unknown how the values might vary across the genome, we consider a simpler system with uniform birth probability, constant death probability, and a fixed number of LEFs. In this formulation, LEF dynamics are well descried by the LEF ***processivity*** (2/death rate, since each head of a non-stalled LEF translocates by 1 lattice-site per time-step), and ***separation*** (total number of lattice sites / number of LEFs). Processivity is additionally interpretable as the average size of a loop extruded by an unobstructed LEF over its lifetime. Small elaborations of this basic 1D model can allow for LEF pausing, permanent stalling at particular genomic elements, active loading and unloading, or even multiple classes of LEFs with different processivity. Note that while 1D dynamics of LEFs are taken as independent from the 3D dynamics of the polymer chain, the interactions induced by LEFs cannot simply be described as an effective pairwise potential. This is because the potential of interactions created by a loop-extruding factor is time-dependent, and LEFs are not independent as they can block each other while on DNA.

### 3D simulations

To perform Langevin dynamics polymer simulations we used OpenMM, a high-performance GPU-assisted molecular dynamics API (Eastman and Pande, 2010; Eastman et al., 2013). To represent chromatin fibers as polymers, we used a sequence of spherical monomers of 1 unit of length in diameter. Here and below all distances are measured in monomer sizes (∼3 nucleosomes, ∼10nm), density is measured in particles per cubic unit, and energies are measured in kT.

Neighboring monomers are connected by harmonic bonds, with a potential *U* = 100(*r* – 1)^2^ (here and below in units of kT). Polymer ***stiffness*** is modeled with a three point interaction term, with the potential *U* = *S* (1 – cos(*α*)), where alpha is the angle between neighboring bonds, and S is a stiffness parameter.

To allow chain passing, which represents activity of topoisomerase II, we used a soft-core potential for interactions between monomers, similar to (Le et al., 2013; Naumova et al., 2013). All monomers interacted via a repulsive potential

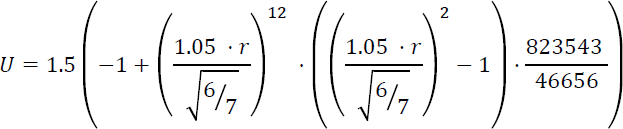

This is a fast and efficient potential designed to be a constant of 1.5 kT up to r=0.7-0.8, and then quickly go to zero at r=1.05.

Simulations were performed in periodic boundary conditions, with the size of the cubic box set to achieve spatial ***density*** of 0.05 or 0.2.

To connect 1D LEF simulations with 3D polymer simulations, we first run 1D LEF dynamics for 4 million time steps to achieve thorough equilibration of LEF positions even in most dense regimes. We then assigned bonds to the monomers in polymer simulations according to the current position of LEFs. The two monomers held by the two heads of each LEF were connected by a harmonic bond with the potential *U* = 4*r*^2^. We then performed a certain number of steps of Langevin dynamics, defined by a parameter setting the number of 3D-simulation time-steps per 1D-simulation time-steps (***3D-to-1D dynamics*** parameter). After that, we advanced LEF dynamics and restarted polymer simulations from the final position of the previous run, but with new position of LEFs (i.e. bonds). Since shifting bond positions in OpenMM is impossible without a computationally expensive re-initialization procedure, for computational efficiency, we advanced LEF dynamics by four steps at a time.

We note that this coarse-grained description of extrusion does not necessarily assume the chromatin fiber is threaded through the LEF, and the processes we describe could be realized by anything that acts processively along the chromosome to generate progressively larger loops.

We performed simulations of a polymer chain consisting of 8 groups of three TADs of 300, 600, and 1200 monomers each, arranged sequentially (300, 600, 1200, 300, 600, 1200…), totaling (300 + 600 + 1200) * 8 monomers. Impermeable BEs were placed between each pair of neighboring TADs. Simulations were started from a compact polymer conformation, as described in (Imakaev et al., 2015), created on a cubic lattice a box of the size (PBC box - 2). Simulations were then advanced for 100 blocks of simulations (400 steps of LEF dynamics). After that, 500 blocks of simulations were performed and their conformations were recorded. After that, LEF dynamics was advanced by 4 million steps, and the process was repeated (100 blocks of simulations, then 500 recorded blocks of simulations) three more times. The latter was done to achieve better averaging. Overall, this yielded 2000 conformations for each parameter set.

To achieve better averaging of contact maps (which are more difficult to average than *P(s)* curves (Naumova et al., 2013)) for all main and supplemental figures, the above numbers were increased as follows: 200 non-recorded blocks of simulations, 1000 blocks of simulations recorded, repeated 10 times to yield 10000 conformations. This changed the goodness-of-fit by no more than 0.03, showing that our parameter sweep produced sufficient number of conformations to evaluate *P(s)* curves.

### Generating simulated contact maps

Generating contact frequency maps for comparison to Hi-C data relies on a final parameter, the ***Hi-C capture radius,*** which is the distance at which two monomers are determined to be in contact. Contact maps were generated using different values of contact radius: 2, 3, 4, 5, 6, 7, 8, 10. To generate a contact map, each conformation was separated into 7 blocks of two TADs (TADs 0,1; 1,2; 2,3… 6,7). Then for each block of two TADs, a contact map was calculated. Resulting 7*2000 contact maps from each of the 2000 conformation were summed together and saved.

To calculate *P(s)* plots from simulated data, a contact map with 1 monomer resolution was used. The contact map was not normalized, since the final *P(s)* curves were allowed to be vertically shifted on a log-log plot (i.e. multiplied by a constant) for calculating goodness-of-fit.

For display, simulated contact maps were first binned using 4 monomer (2.4 kb) bins. Then, the map was normalized such that an average sum over all rows equals one. The map was then clipped at 0.0035; values of zero were shown as white, values of 0.0035 or larger were shown as black.

### Parameter sweep

To investigate the features of simulated contact maps, we varied the above parameters over a wide range of values. In particular, we varied 5 simulation parameters:

LEF processivity: 100, 200, 400, 800, 1200, 1600
LEF separation: 50, 100, 200, 400, 800, 1600
3D-to-1D dynamics: 300/4, 1000/4, 5000/4
Stiffness S: 0, 2, 4, 6
Density: 0.05, 0.2

For each of these 864 separate simulations, we generated 2000 conformations (total 1,728,000). From these, we calculated contact maps for 8 Hi-C capture radius values: 2, 3, 4, 5, 6, 7, 8, 10. Together, this gave 6912 parameters sets.

### Simulated FISH CDFs

For each parameter value, simulated FISH CDFs were calculated by aggregating distances for for a pair of ‘within-TAD’ loci in the largest (1200 monomer) TAD and a pair of ‘between-TAD’ loci that span the BE between the 600 and 1200 monomer TAD, both at a separation of 600 monomers (360kb) for the same sets of conformations used to build contact maps.

### Simulated P(s)

From each contact map, we calculated *P(s)* as follows. For each diagonal of the contact matrix, we evaluated the average values of contact probability within 300-monomer TADs, 600-monomer TADs, 1200-monomer TADs and regions not belonging to any TAD (trans). We then averaged the values in logarithmically-spaced bins with a step of 1.2 starting at 7 (7, 8, 10, 12, 15, …,x,x*1.2, x*1.2^2,…, N) where N is the length of a given *P(s)* curve (300, 600, 1200 for TADs, 1300 for between-TADs, in monomers).

### Experimental P(s)

To calculate experimental *P(s),* we used publicly-available data from (Rao, 2014 (Rao et al., 2014)), processed in-house using ICE (Imakaev et al., 2012), for GM12878 inSitu protocol and Mbol restriction enzyme. Data was binned at 2kb. Since a whole-chromosome 2kb resolution contact map is too large to fit in memory, we split the data in blocks of 20Mb, overlapping by 10Mb (starting at 0, 10Mb, 20Mb, 30Mb, etc.). We removed bins with <100 reads, and then iteratively corrected each block.

We obtained TAD annotation from (Rao, 2014), filename: GSE63525_GM12878_primary+replicate_HiCCUPS_looplist.txt.gz. For all annotated TADs, we found a 20MB block most centered at this TAD, and evaluated P(s) within this TAD. To calculate *P(s)* between TADs, we defined a set of boundaries as the union of start and end-points for all TADs. P(s) was then calculated between loci-pairs upstream and downstream of the boundary up Mb separation.

### Experimental Hi-C maps

We used the same dataset to display regions of the Hi-C map in **Fig 1** and **fig. S1.** Datasets were binned at 5kb resolution to reduce sampling noise. Data were corrected in the same way as for *P(s)* calculations.

### Goodness-of-fit

To compare experimental and simulated *P(s),* we first selected experimental TADs that are comparable in size to each simulated TAD. We selected TADs that are between 0.9 and 1.1 of the size of simulated TAD, assuming each monomer is 600kb (giving 180kb, 360kb and 720kb TADs). We then calculated median *P(s)* for TADs in each category, as well as 10^th^ and 90^th^ percentiles. *P(s)* was calculated in logarithmically spaced bins, as described above. The three curves were plotted for comparison with simulated scalings. Next, we evaluated experimental *P(s)* at the same values of *s* as simulated *P(s)* curves were calculated. Note that since these values are logarithmically spaced, each portion of the log-log plots shown contribute equally. We then define the goodness-of-fit as the *geometric standard deviation* of the ratio of simulated to experimental *P(s)*; this has an interpretation as a typical fold-deviation of *P(s).*

A value of 1 indicates perfect fit. Larger values would indicate worse fits. The best fit we observed was 1.103 (processivity 200, separation 200, 5000 steps, stiffness 2, density 0.2, cutoff 10). The second best fit was the same as first but with processivity of 400 (fit 1.110). The worst fit had a value of 4.46 for (processivity 100, separation 50, 1000 steps, stiffness 0, density 0.05, and cutoff 2). The poor fit in this case is likely because the small tight loops formed in this regime formed a very dense fiber that has many fewer long-range contacts than experimental data. Notably, the second worst fit was similar, while the third worst fit, 4.03, had completely different parameter values: (processivity 100, separation 1600, 5000 steps, stiffness 0, density 0.05, and cutoff 2), which lies in the free polymer regime. All parameter values and fit coefficients are summarized in **Database D1.**

Since the best fitting models had diverse parameter values, we took first 100 best fitting models (fit values 1.103 to 1.195) and assessed how frequently each pair of (processivity, separation) occurs in this list, and what the best fit was for each pair (see **Fig 2B**).

We note that any biological realization of active linear compaction by loop extrusion has many additional unknowns beyond the parameters we use to describe the process; this can include the size of the LEF and the interactions between LEFs and other DNA-bound proteins. The former may affect both the distance between LEF proteins at which they become stalled, as well as the distance between two regions of DNA bound by a LEF. Moreover, experimentally-mapped Hi-C TADs display a great deal of diversity (**fig. S1**), and would most likely be described by different sets of parameters. For these reasons, we focused on which combinations of parameters frequently gave a good fit, and the characteristics of TADs in this parameter regime, rather than relying on particular best-fitting parameter set. We found that while nearly any value of any parameter can be found in the top parameter-sets, we find that certain pairs of LEF processivity and LEF separation are found more often.

### Modifications of the main model

To simulate a model of semi-permeable boundary, we introduced a genomic element in the middle of the longest TAD that permanently stalls LEF heads with a probability of 20%. We note that the effective stall rate is actually higher, as once one LEF becomes stalled at the boundary, another LEF is likely to stall against the original LEF, etc. Thus, only a small probability to stall LEFs is needed to “nucleate” a TAD boundary, even in a relatively dilute regime. In our case, a probability of 0.2 creates a strong visible TAD boundary.

To create a model with uneven loading of LEFs, we increased the birth rate 30-fold in the region of the length of (TAD length / 30), located starting at the 10^th^ monomer of the TAD. This made it such that ½ of all LEFs are born next to a TAD boundary, while the other ½ are born randomly.

To simulate folding of the “vermicelli” chromosomes, we increased the processivity in the best-fitting model 10-fold and 20-fold, to 2000 and 4000 respectively. For each value, we observe prominent folding of the chromosome in a linearly organized “vermicelli” state, reminiscent of the first stage of the two-stage model (Naumova et al., 2013) presented in (Naumova, 2015).

### Assigning CTCF motifs to peaks

Motifs were assigned to narrow peak calls by interval intersection using bedtools (Quinlan and Hall, 2010). If more than one motif mapped to a single peak (0.5% of hits), only the first motif found was assigned.

### ChIP-seq peaks around oriented CTCF motifs

The CTCF peaks identified in GM12878 (GSM935611) that have an overlapping motif occurrence (PFM: CTCF_known1) from (Kheradpour and Kellis, 2014) were ordered by fold enrichment value. The directionality profiles are centered at the peaks, but oriented so that all motif instances “point” in the 3’ direction. We then selected the 4000 most and least enriched CTCF peaks from this set and produced histograms of the summit positions of all called ChIP peaks for CTCF (GSM935611), Smc3 (GSM935376) and Rad21 (GSM935332). Histograms are centered at the peaks, but oriented so that all motif instances “point” in the 3’ direction.

### Model of orientation-specific BEs with varying permeability

To convert ChIP-seq peak strength to the occupancy of simulated BEs, we performed the following steps: first, we calculated the middle of each ChIP-seq narrow peak as the (start + end)/2. The orientation of each CTCF peak was then assigned using the distance to the nearest rad21 peak, as not all CTCF peaks have a detected CTCF_known1 motif. Separately for each orientation, we then calculated a binned profile of the sum of peak fold-change-over-input values for peaks whose centers fall in each 600bp bin. Bins with no peaks were assigned a value of -infinity. We then transformed these values with a logistic function f(x)=1/(1 + exp(-x/20 - mu)), where mu was selected to be 3, and changed to 2 or 4 to respectively impose a greater or less amount of bound CTCF, and 20 was selected to allow for a wide range of BE occupancies. We used a logistic function, as it constitutes a natural way to convert from arbitrarily dispersed values to numbers between 0 and 1. The resulting profiles were used as BE permeability for each orientation.

We modelled a 15MB region of human chr14, 60,000,000 to 75,000,000, which was chosen as it has gene deserts, gene-rich regions, and is uninterrupted by poor coverage bins. Simulations were performed in two replicates for each region, and 50,000 conformations (steps of SMC dynamics) were obtained for each replicate. The resulting contact map was then calculated, and averaged in 20×20 blocks, corresponding to 12kb resolution, to obtain a 1250x1250 contact map.

To compare simulated contact map with the Hi-C contact map, we used GM12878 *in Situ* data from (Rao et al., 2014), mapped, filtered and binned using *mirnylib* software for each restriction enzyme (MboI and DpnII) and then pooled together. The data was obtained at 10kb resolution, linearly interpolated to 6kb resolution, and block-averaged in 2×2 blocks to obtain a 12kb-resolution contact map (1250x1250 bins).

To calculate spearman correlations between experimental and simulated contact maps as a function of genomic position, we first removed the strong effect of genomic distance on interaction frequency; as previously(Naumova et al., 2013) the observed interaction matrices were divided by an expected interaction matrix, calculated as the mean number of interactions at a given distance, using a sliding window with a linearly increasing size. Spearman correlations were then calculated using sliding a genomic window of 3Mb along by 1Mb maximum separation.

### Model of direct BE-to-BE interactions

For simulations with direct BE-to-BE interactions, the inter-monomer interaction energy was calculated for r < attractionRadius, and was zero otherwise. The expression below defines U, the energy of interaciton between any two monomers. Sticky1 and Sticky2 are index variables that are equal 1 for sticky BEs, and zero for all other monomers. Function step(x) is a step-function; equals 1 for x>0, and zero otherwise. attrE is attraction energy in kT, was selected to be 1.5, 3, and 5. r is distance between centers of the particles.

emin12 = 46656.0 / 823543.0; rmin12 = sqrt(6.0 / 7.0)); attractionRadius = 1.5; repulsionRadius = 1; ATTRdelta = (attractionRadius - repulsionRadius) / 2.0); rshft = (r - repulsionRadius - ATTRdelta) / ATTRdelta * rmin12; ATTReTot = min(Sticky1, Sticky2) * attrE * kT;

Eattr = - rshft^12 * (rshft^2 - 1.0) * ATTReTot / emin12 - ATTReTot; rsc = r * rmin12;

Erep = (rsc^12 * (rsc^2 - 1.0) / emin12 + 1) *3*kT; U = step(1 - r) * Erep + step(r - 1) * Eattr;

This function is described in our bitbucket repository: https://bitbucket.org/mirnylab/openmm-polymer, revision 9b4303b from 2015-12-27, file openmmlib/openmmlib.py, line 1213. Simulations were performed in 10 independent runs, recording 1000 blocks of 10000 MD steps each after skipping the first non-recorded 100 blocks. Contact maps were displayed for contact radius 5 and 10.

### Models of bulky and stiff boundary elements

To simulate a of bulky BEs, we started with the same polymer chain of 8 groups of three TADs of length 300, 600 and, 1200 monomers as used in the models with loop extrusion. However, instead of introducing any LEFs, several polymer chains were connected at each BE (either 3 of length 10 attached 1-per-monomer to the monomers around the BE via harmonic bonds, or 5 chains of length 6). For stiff BEs, the 10 monomers around the BE had an increased stiffness of 6, while other monomers had a stiffness of 1, as defined above. As there were no LEF dynamics, conformations were recorded every 50000 MD steps, after an initial period of 100 non-recorded blocks for a total of 1000 blocks and contact maps were averaged over 5 independent runs. Contact maps were displayed for contact radius 5 and 10, as other values produced similar results.

---

**Fig. S1.**
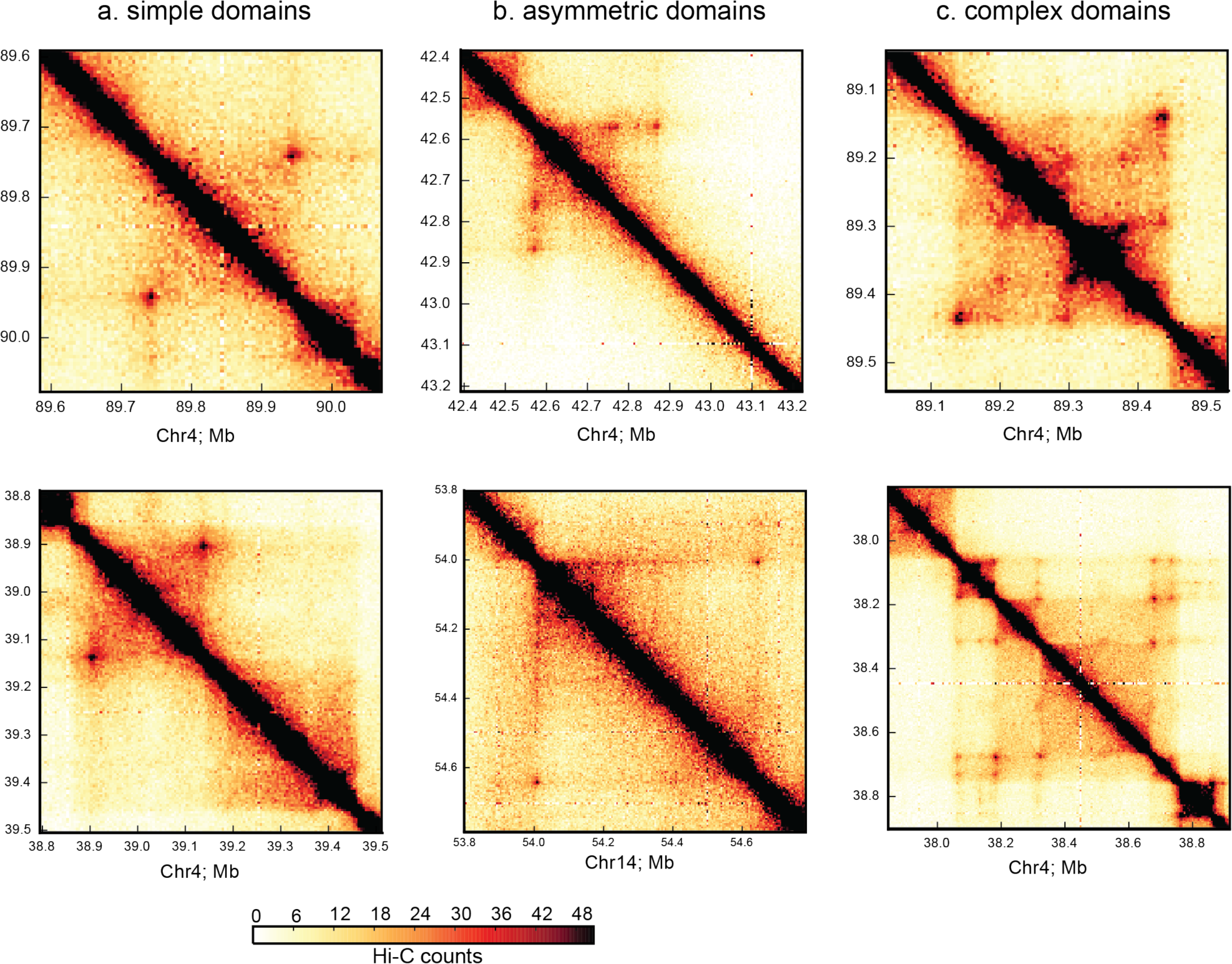
Diversity of TADs. Experimental Hi-C contact maps at 5kb resolution for GM12878 in-situ (Rao et al., 2014), showing: **a.** simple, **b.** asymmetric, and **c.** complex TADs. Note a TAD without a corner-peak in the bottom left figure.

**Fig. S2.**
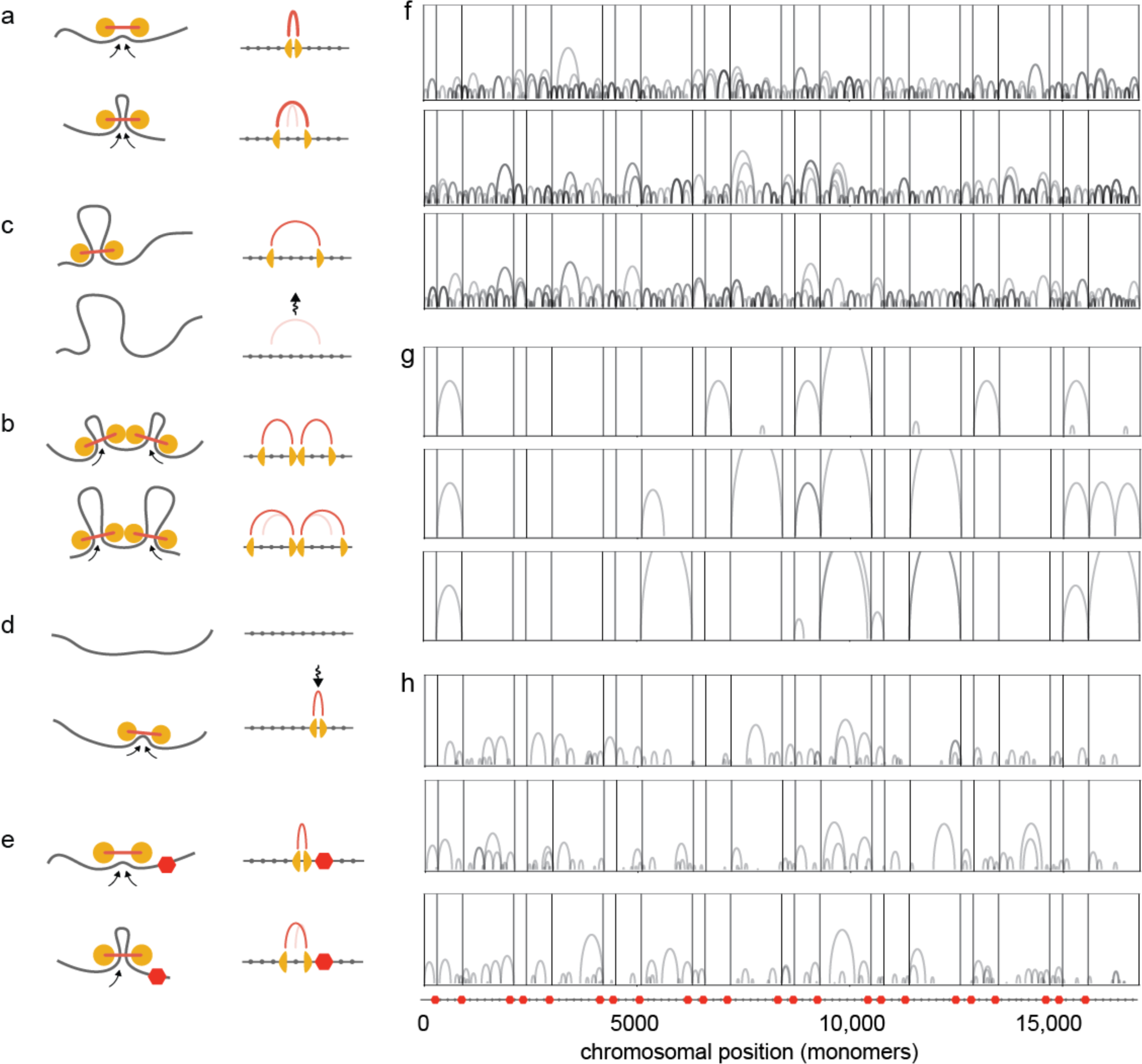
Arc-diagrams of loop interactions imposed by LEFs. **a-e.** Illustration of LEF update rules and lattice used for simulation of 1D LEF dynamics. As in the main figure, LEFs shown as linked pairs of yellow circles, chromatin fiber in grey. In the associated 1D lattices, LEF positions are shown by yellow triangles, and red arcs indicate the lattice sites (monomers) that are held by the LEF before and after the update. **a.** extrusion, **b.** stalling, **c.** dissociation, **d.** association, **e.** stalling at a BE (red hexagon); cartoons shown next to and their associated 1D lattice updates. **f-h.** Three examples shown to highlight the diversity of looping interactions for three processivity-separation. Arcs indicate monomers held together by a given LEF. Arc height reflects loop size, and reinforced loops (i.e. those held by multiple LEFs) are in darker shades of grey. Positions of boundary elements (BEs) shown by vertical bars, illustrating how loops only form in-between neighboring BEs. Lattice with position of BEs as red hexagons shown schematically below. **f.** processivity 720kb, separation 30kb. In the limit of small separation and large processivity, LEFs form a densely packed array of consecutive nested non-overlapping loops stretching between consecutive BEs. **g.** processivity 960kb, separation 960kb. In the limit of large processivity and large LEF separation, LEFs extrude the entire region between, and are stably bound at, consecutive BEs, forming a loop between the two boundary elements with high probability. **h.** processivity 120kb, separation 120kb. In this regime of medium processivity and medium separations, both 2-4 times less that a TAD size, LEF dynamics encompass many events: a small fraction of LEFs happen to live long enough to extrude the entire region and transiently connect two BEs, although most LEFs dissociate before reaching the nearest boundary.

**Fig. S3.**
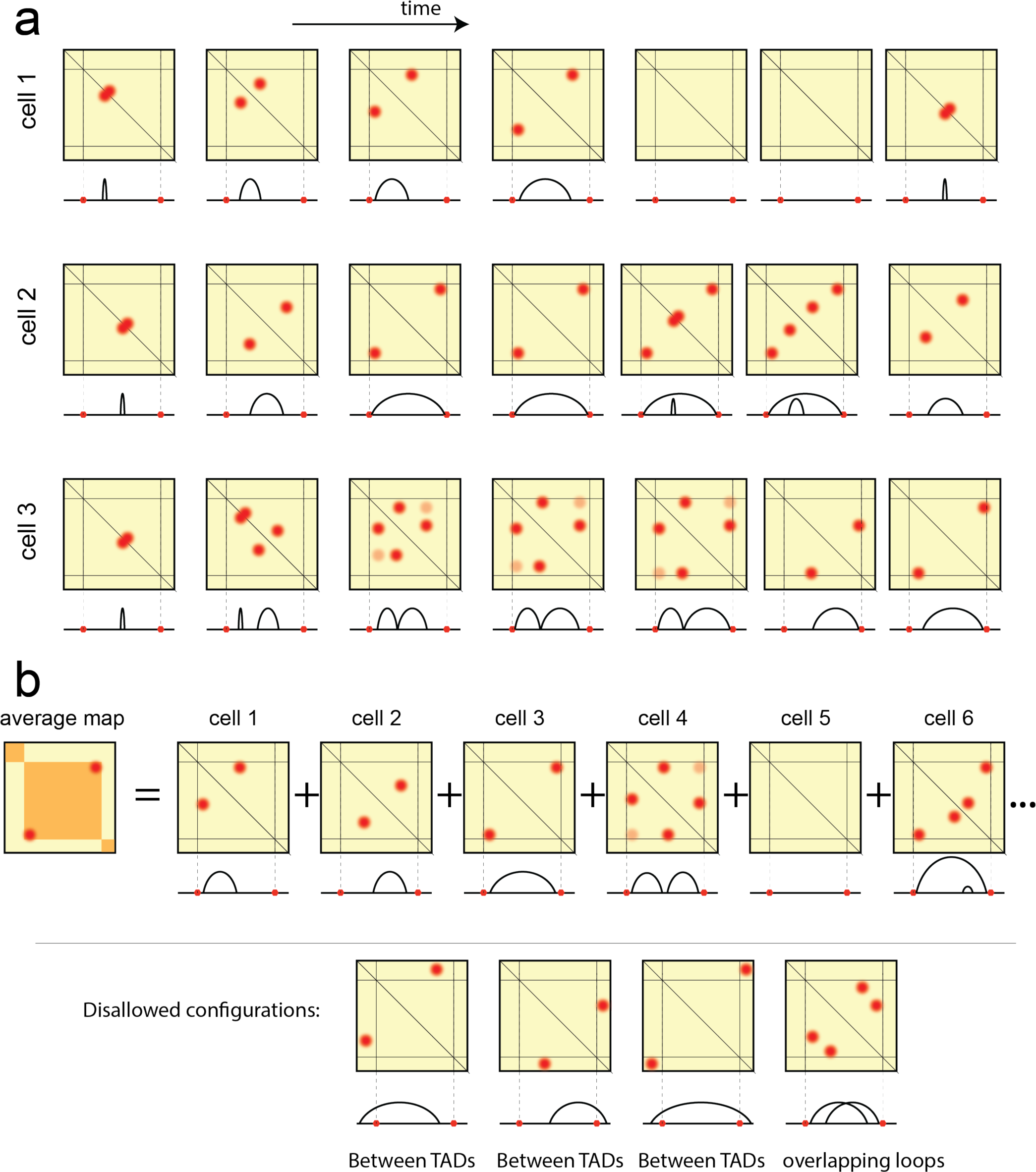
Illustration of LEF dynamics. **a.** Loop extrusion is a stochastic process that follows a different trajectory in each cell. This figure outlines possible dynamics of LEFs using three examples. Each trajectory consists of 7 snapshots (frames), showing LEF positions as arches, and LEF-mediated loops (red dots) on the schematic single-cell Hi-C map. BE positions are indicated as red hexagons on the arch diagram, and as lines on the Hi-C map. For cell 1, a LEF associates to the chromatin fiber (frame 1), and extrudes a loop until it meets a boundary (frame 3). It then continues to extrude a loop from the other side (frame 4), but soon dissociates (frame 5). After some time, a new LEF lands in a different location (frame 7). For cell 2, a LEF associates to the chromatin fiber (frame 1) and continues to extrude a loop until both sides of a LEF are stalled against boundaries (frame 3). A LEF stays in the same stalled position (frame 4), while a new LEF associates to the chromatin within the loop created by the original LEF (frame 5). This new LEF continues to translocate, and the old LEF stays (frame 6), and finally dissociates (frame 7). In cell 3, a LEF associates to the chromatin (frame 1), and another LEF associates nearby (frame 2). The two LEFs stall against each other (frame 3), which creates a weaker interaction between the beginning of the first LEF and the end of the second LEF as they are now nearby in space. The two LEFs continue to translocate until each of them hits a BE (frames 4 and 5). The first LEF then dissociates (frame 6), and the second LEF starts to translocate again (frame 7). This example illustrates how corner-peaks may occur at distances longer than the LEF processivity, because they can be mediated by a chain of two or more LEFs stalled against each other. **b.** Hi-C maps are formed as an average over many cells. In each of the cells, the configuration of LEFs is different. However, in each configuration LEFs only form loops within TADs (i.e. not crossing BEs). Note that because LEFs stall at BEs, many configurations will have a LEF connecting two ends of a TAD. As a result, an average map may have a peak of interactions between a start and an end of a TAD. We also note that direct BE-to-BE loops are not the only contributors to the peak of interactions between neighboring BEs. As shown in cell 4, two LEFs stalled against each other and against neighboring BEs will bring the BEs close together in space, and thus contribute to the peak of interactions. In the more dense regimes of LEF dynamics (small separation, high processivity), even longer consecutive LEF arrays may contribute to the BE-to-BE interaction peak. Certain configurations of loops are disallowed in the LEF dynamics described in this paper. In particular, a LEF will never connect two regions separated by a BE. Similarly, two loops will never cross (though they could be nested, as shown above). These examples are shown in the bottom of the figure.

**Fig. S4.**
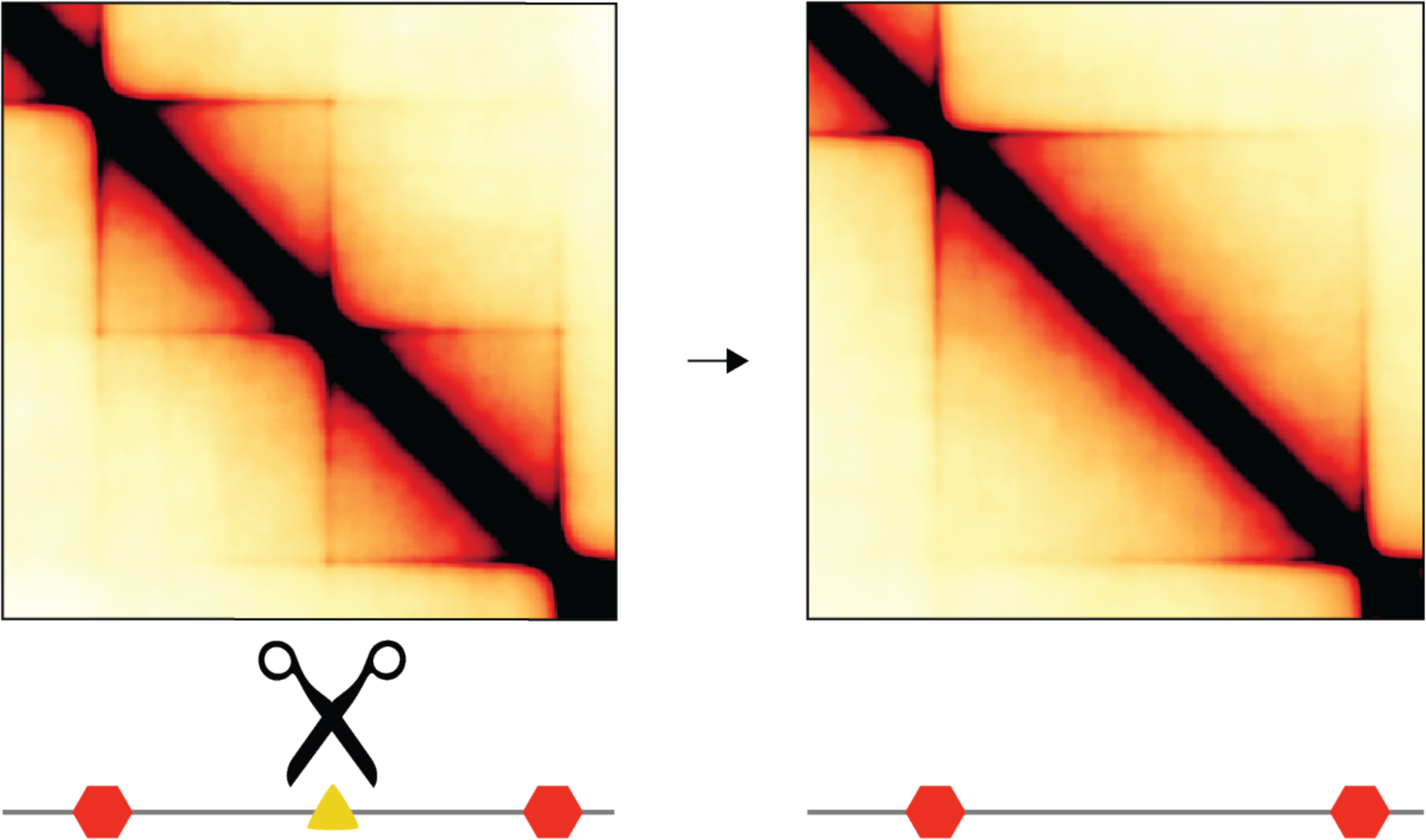
Simulated semi-permeable boundary deletion. Simulated deletion of a 50%-permeable boundary element (as compared with 10% in **Fig 5**) halfway between two boundary elements separated by 720kb.

**Fig. S5.**
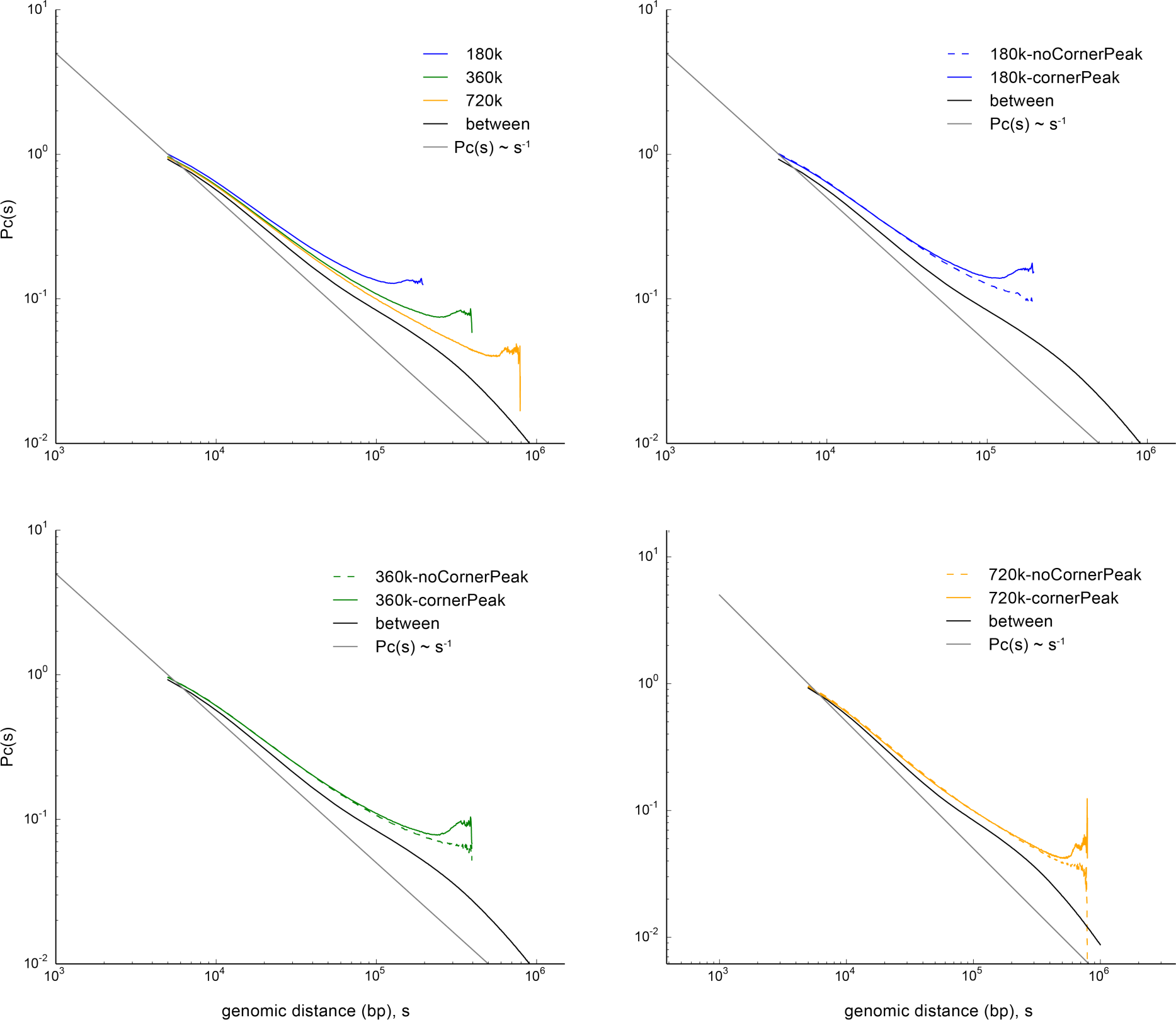
Average contact probability vs. genomic distance. Average contact probability, *P(s)*, vs. genomic distance, *s*. Calculated from 2kb resolution GM12878 in-situ DpnII maps (Rao et al., 2014). *Top left*: For TADs of 180kb, 360kb, and 720kb, and between TADs using published TADs (GSE63525_GM12878_primary+replicate_Arrowhead_domainlist.txt.gz (Rao et al., 2014)). *Top right*: For TADs of 180kb with and without corner peaks, as determined by having a published peak less than 50kb from the TAD corner (sum of distances between starts and between ends of peak/TAD). Published peaks from GEO database, GSE63525_GM12878_primary+replicate_HiCCUPS_looplist.txt.gz (Rao et al., 2014). *Bottom left*: similarly, for TADs of 360kb. *Bottom right*: similarly for TADs of 720kb.

**Fig. S6.**
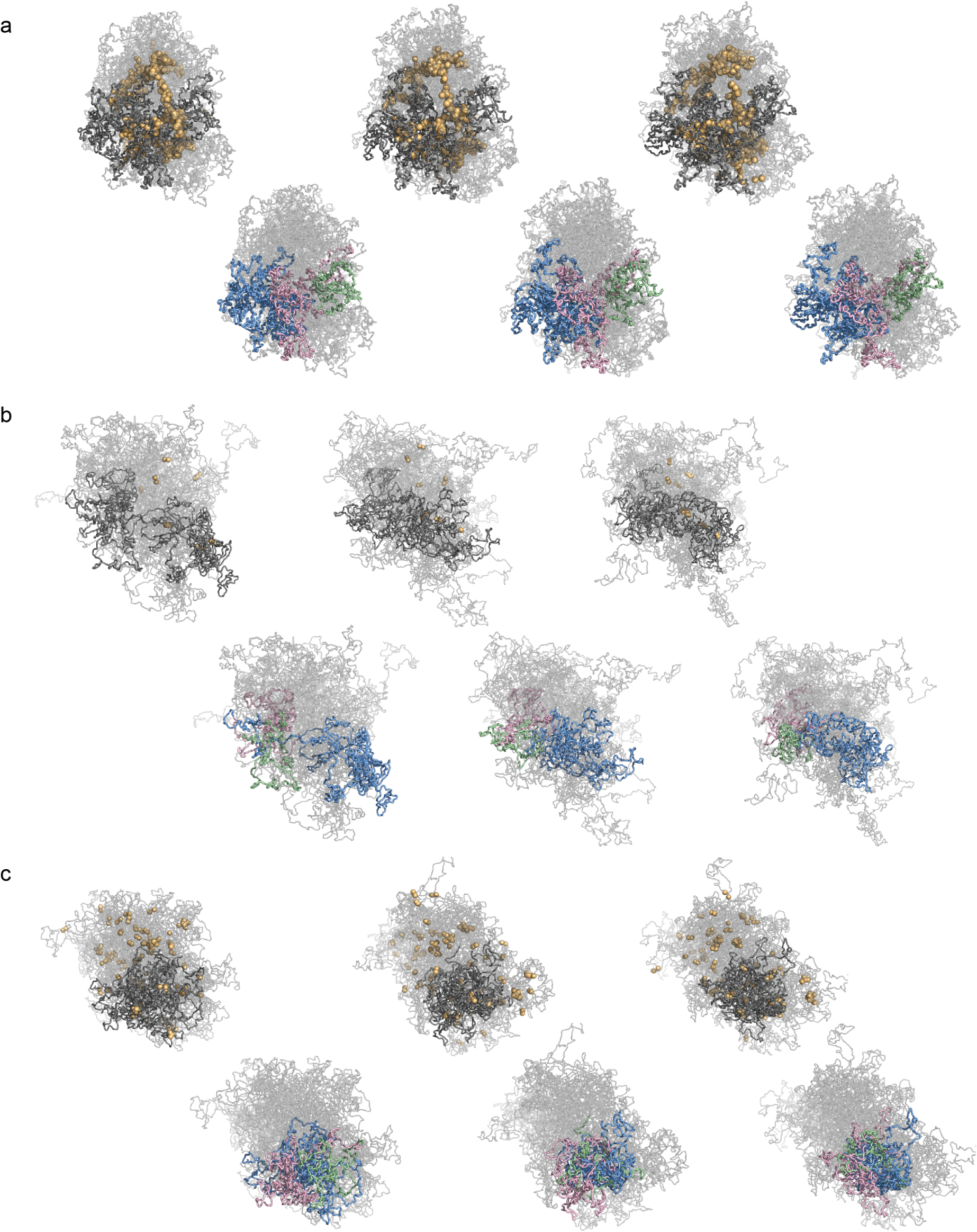
Spatial properties of TADs organized by LEFs. Conformations of a polymer subject to LEF dynamics for three nearby time-points. For each, the top row shows LEFs (yellow spheres), and chromatin (grey), for three conformations; bottom row shows three neighboring regions between BEs of sizes (180kb, 360kb, 720kb) in (green, pink, blue) for the same conformations. **a.** processivity 720kb, separation 30kb. **b.** processivity 960kb, separation 960kb. **c.** processivity *120kb*, and separation 120kb (as in **Fig. 1D**).

**Fig. S7.**
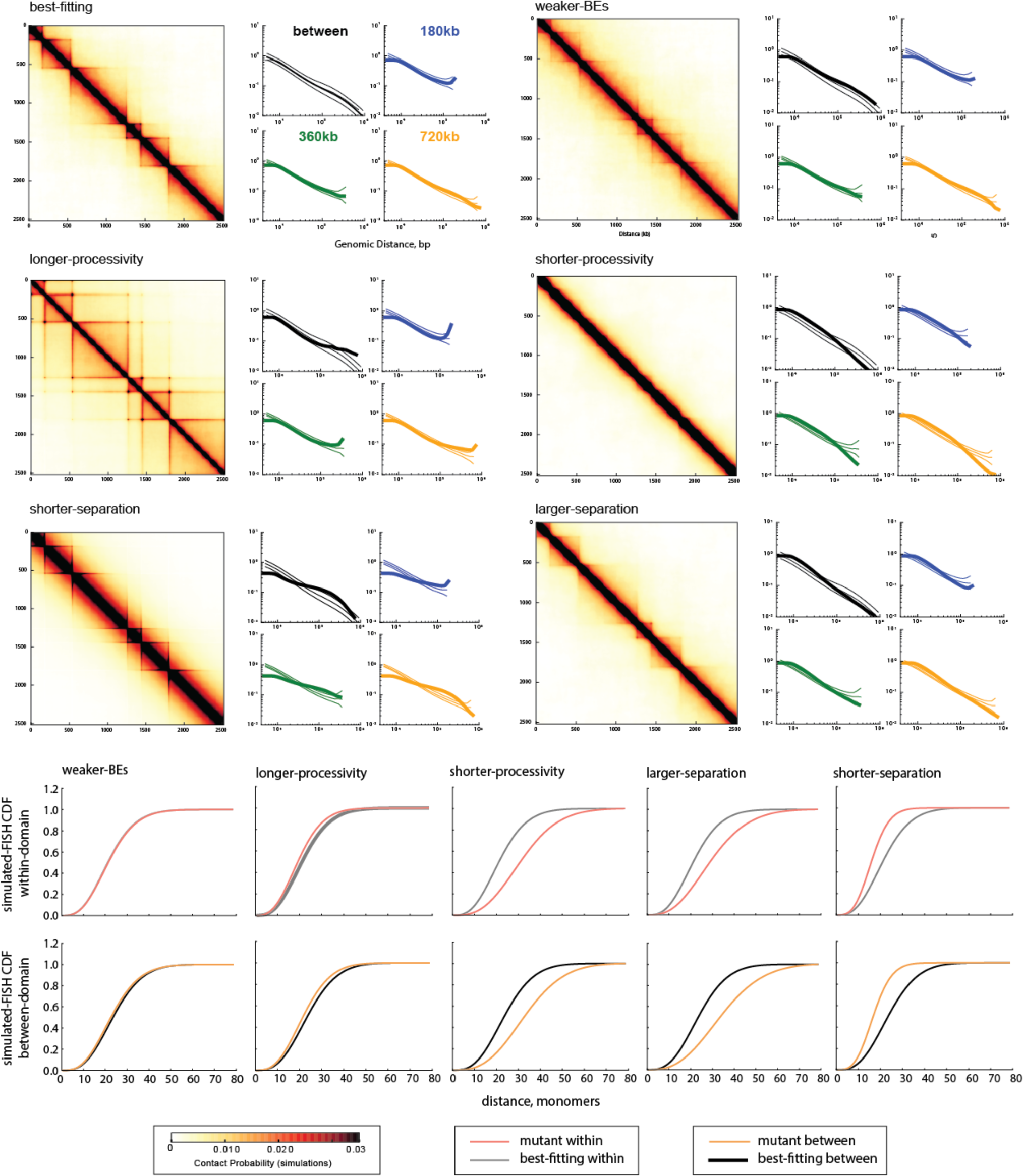
Experimentally testable predictions from pertubrations of LEF dynamics. ***Top***: Contact maps and P(s) for perturbations of the best-fitting model (processivity 120kb, separation 120kb, 5000 steps, stiffness 2, density 0.2, cutoff 10.) ***Bottom***: Simulated FISH CDF functions for pairs of region within the TAD, and in two neighboring TADs, both separated by 600 monomers (360 kb). Grey or black line corresponds to the best-fitting model, while orange line corresponds to the perturbation of the model. Perturbations of the model are as follows: *For weaker-BEs*, all BEs were made semi-permable, with 50% permeability. For *longer-processivity, shorter-processivity, shorter-separation*, and *larger-separation* models, the processivity or separation were respectively increased or decreased by a factor of 4 compared to the best-fitting model. The models correspond to possible experimental perturbations: *Weaker-BEs* can correspond to CTCF depletion. *Longer-processivity* may correspond to the decreased unloading of LEFs, which can be achieved experimentally by depleting the LEF unloader (e.g. for cohesin, Wapl); *shorter-processivity* may come from over-expressing the unloader. *Shorter-separation* may be achieved by increasing the number of chromatin-bound active LEFs, either by increasing loading of LEFs (e.g. for cohesin, Nibl), or by overexpressing LEFs; *larger-separation* can come from the opposite perturbations. Results of the perturbations are the following: *Weaker-BEs* (CTCF depletion) shows blurred TAD boundaries on the contact map. The FISH CDF doesn’t change much for within-TAD probes, but the between-TAD CDF shifts to be closer to the within-TAD CDF. *Higher-processivity* displays very strong corner peaks. Spatial distances decrease slightly, but not as much as in other processivity or separation changes. *Shorter-processivity* model displays an almost complete loss of TADs, has increased distances in FISH for both within- and between-TAD, and makes the two curves more alike. *Shorter-separation* tilts *P(s)* curves to be more steep and decreases FISH distances. It also leads to sharper TAD boundaries. *Larger-separation* makes TADs less pronounced and leads to an increase in FISH distances within and between TADs.

**Fig. S8.**
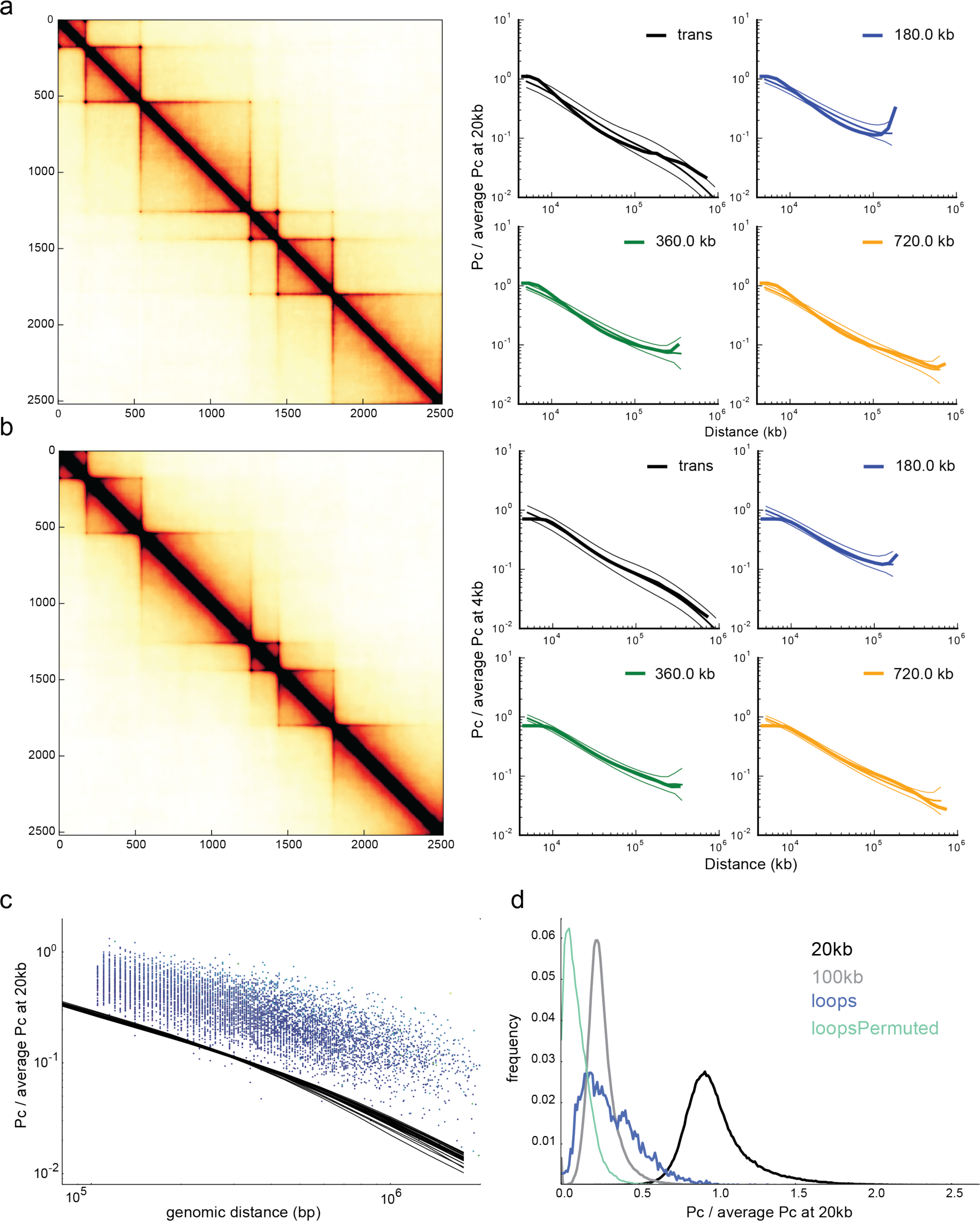
Corner-peaks are transient and naturally emerge at higher LEF lifetimes. **a.** *Left*: contact map showing TADs and peaks, with processivity 240kb, separation 120kb. *Right*: contact probability vs. distance for TADs of three sizes, and between-TADs (trans). Corner-peaks are connected by a LEF (25%, 7%, 0.2%) of the time for TADs of (180kb, 360kb, 720kb). **b.** as above, but for processivity 120kb, separation 120kb, as in **Fig 1E.** *left.* Corner-peaks are connected by a LEF (9%, 0.7%, <0.01%) of the time for TADs of (180kb, 360kb, 720kb). Together, **a** and **b** show increasing LEF processivity naturally strengthens peaks at TAD corners. Note that for the increased processivity, the corner-peak for the largest 720kb TAD is clearly visible on the contact map and gives a slight increase of *P(s)* at the end of the plot. Yet, the start and the end of this TAD are connected by a LEF only 0.2% of the time. This emphasizes that corner-peaks are not permanent loops; permanent loop would have much more drastic effect on the contact map. **c.** contact probability *P(s)* vs. genomic distance for corner-peaks (blue circles) using published locations from GM12878 and the associated Hi-C contact maps at 10kb resolution, normalized to 1 at 20kb. Black lines show average chromosome-wide *P(s)* (one line per chromosome). **d.** histogram of contact probability for all published peak-pairs (blue) versus all loci separated by 20kb (black) or by 100kb (grey), and permuted peak-pairs (green, with shuffling along diagonals of the Hi-C map to preserve the distribution of peak-pair genomic distances). Together, **c** and **d** show that peak-loci do not appear to be in permanent contact in Hi-C data, and corner-peaks are best considered as transient loops.

**Fig. S9.**
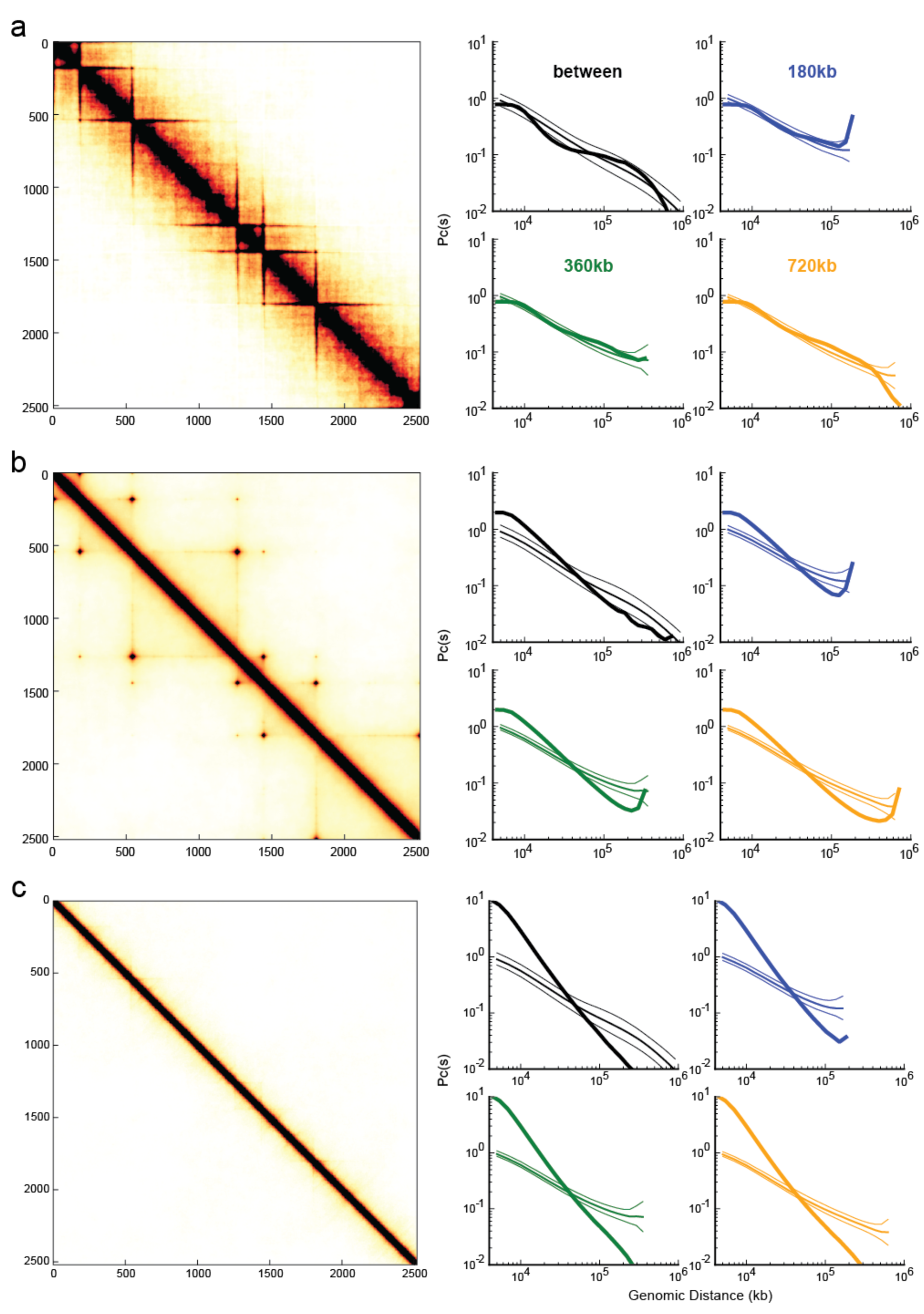
Diversity of regimes from LEF dynamics. *Left*: contact maps. *Right*: associated *P(s)* for three TAD sizes and between TADs. **a.** An example of a dense regime that results in a contact map with very strong domain borders, yet fits the scaling quite well and is found within 100 best-fitting models: goodness-of-fit of 1.1828 (rank 65). Parameters of the model are: processivity 720kb, separation 30kb, 3D-to-1D ratio 300/4, stiffness 0, density 0.05, cutoff 8. **b.** An example of a “simple loop” regime with poor fit of 1.4137 (rank 2208 out of 6912). Processivity 960kb, separation 960kb, 3D-to-1D ratio 1000/4, stiffness 2, density 0.2, cutoff 8. This regime display strong loops between neighboring BEs. This example also shows that simply bringing neighboring BEs together is insufficient to create TAD observed in Hi-C. **c.** In the ‘free-polymer limit’ of short processivity and large separation, the effect of LEFs on the polymer is minimal. *Left*: contact map showing neither TADs nor peaks. *Right*: contact probability vs. distance for TADs of three sizes, and between-TADs (trans). Processivity 60kb, separation 960kb, 3D-to-1D ratio 5000/4, stiffness 0, density 0.05, cutoff 5.

**Fig. S10.**
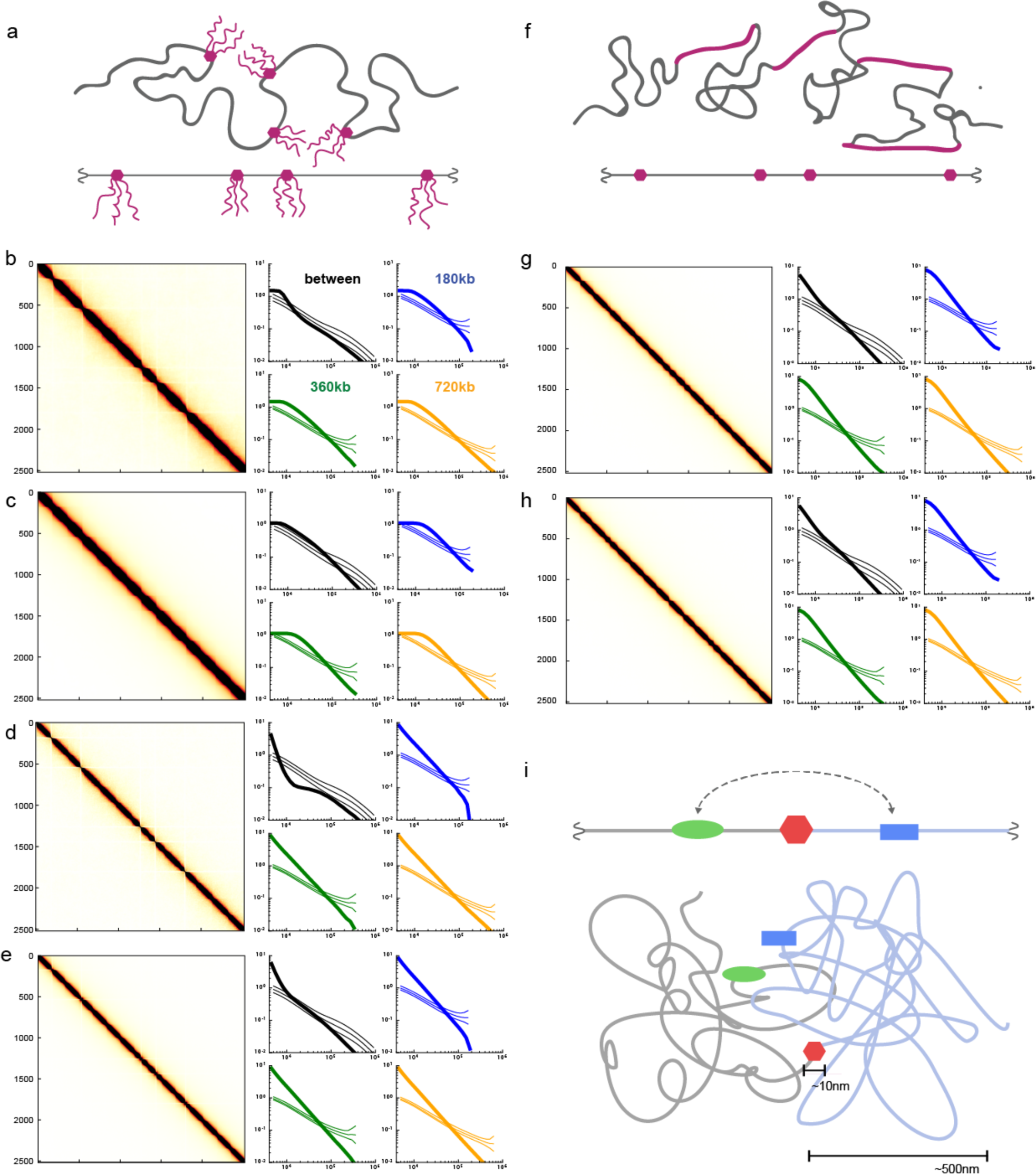
Difficulty of insulating large domains with small genomic elements. **a.** A schematic illustration of a model that explores whether BEs modeled as large bulky objects (e.g. bound by proteins or RNA) can insulate neighboring TADs. In these simulations, each BE is bound by several polymer chains (3 of length 10 in b,d; 5 of length 6 in c,e). Note that since our monomer represents 3 nucleosomes (∼600 kDa), the bound mass would be 18 mDa in each case, which is more than 100 CTCF molecules (82-130 kDa) or 250 Rad21 molecules (72 kDa). This simulation show that insulation between neighboring TADs does not arise from direct physical blocking of interactions between distal genomic regions by BEs. While the bulky BE does create partial insulation between regions directly adjacent to the BE (a “break” in the diagonal of the contact map), this insulation does not extend much further and does not reach the length scale of a full TAD. Moreover, this bulky BE model does not produce peaks of contact probability between proximal BEs. **b-e.** Contact maps and P(s) for a bulky BE model (a) with 3 bristles of length 10 (b,d) or 5 bristles of length 6 (c,e). The contact radius 10 for b,c and 5 for d,e. **f.** A schematic illustration of a model in which we explore whether BEs may correspond to stiff regions of chromatin, thus physically separating two TADs. In this simulation, the majority of the chromatin fiber is flexible (stiffness 1), interspersed with stiff BEs (10 monomers with stiffness 6). While this model also creates insulation between regions directly adjacent to the BE, it does not extend much further and does not reach the length scale of a full TAD. **g, h.** Contact maps and P(s) for the stiff BE model; contact radius 10 for g, and 5 for h. **i**. *Top*: Illustration of a genomic region with an insulating element (red hexagon), a promoter (blue square) and an enhancer (green oval) in 1D. Two neighboring TADs are colored in grey and light blue. *Bottom*: illustration in 3D. Insulation can be easily represented in 1D when the chromosome is drawn as a straight line with the insulating element between the enhancer and promoter. This creates an impression that insulation is relatively easy to achieve. However, intuition developed in 1D is misleading when applied to long flexible chromatin fibers in 3D, particularly because the physical size of an insulating element (3-50nm) is much less than the size of a TAD (300-1000nm). For example, an enhancer and promoter separated by ∼100kb (i.e. much greater than the persistence length of the chromatin fiber) can come in contact while still being spatially far from the insulating element in 3D. Our simulations of BEs as bulky objects or locally very stiff regions argue that it is very difficult to insulate large genomic regions from each other by small factors that act locally. Indeed, our simulations suggest that a crucial mechanistic insight is often overlooked in illustrations of insulating elements. In this work, we propose that insulation at the scale of TADs can be achieved by loop extrusion limited by BEs; crucially, LEFs convert the difficult prospect of insulating in 3D into the more intuitive 1D case and allow insulation to be mediated over spatial and genomic distances much larger than the size of bound proteins forming a BE.

**Fig. S11.**
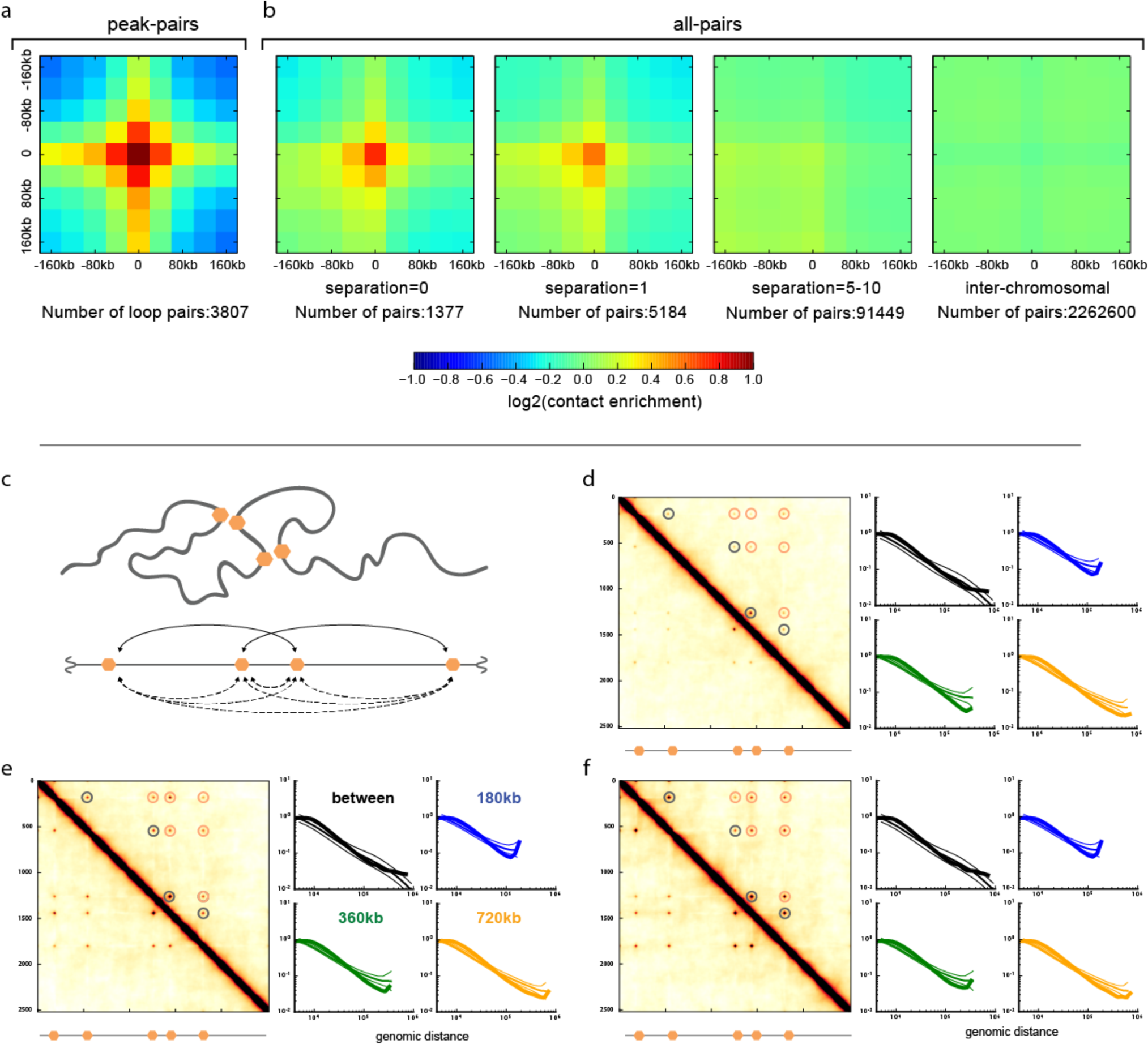
Mediating interactions is not an intrinsic property of peak-loci. **a,b. *Enrichment plots around peak loci*** at 40kb resolution for observed-over-expected GM12878 in-situ (Rao et al., 2014) contact maps (i.e. after removing distance-dependence). **a.** enrichment only around peak pairs called for this dataset (i.e pairs of peak loci, GSE63525_GM12878_primary+replicate_HiCCUPS_looplist.txt.gz, (Rao et al., 2014)). Peak pairs show a pronounced peak in average enrichment. **b.** For this plot, we split the list of all peak pairs into a list of individual unique peak loci. We then calculate enrichment around pairs of peak loci separated by a given number of loci in between. We note that separation of zero is less enriched than that of peak-pairs, as separation zero does not imply that the two peak loci form a corner-peak; they could be the end of one corner peak and the beginning of the next peak. We also calculated enrichments for all between-chromosomal pairs of peak loci. We find that peak loci at further separations display weaker peak in average enrichment. Notably, the peak completely disappears on the between-chromosomal map. This shows that the ability of peak-loci to mediate an interaction peak is not an inherent property, and suggests peaks are realized by a mechanism that acts along the chromosome. We note that a very small (∼1%) depletion of interactions between peak loci between chromosomes, which is barely visible on the interaction map as a bluer cross, is consistent with the idea that loci involved in the formation of peaks are sterically excluded from interactions with other genomic regions (Doyle et al., 2014). **c-f. *models with direct BE-to-BE interactions forms peaks, but not TADs.*** In this alternate model, all pairs of BEs experience a short-range attractive force (with strength 1.5kT, 3kT, and 5kT in d,e,f). When two BEs are far away from each other, they do not interact and move randomly due to the diffusive motion of chromatin. When two BEs come in close proximity (<25 nm), they experience the attractive force (see **Methods** for the attraction energy function). This would happen, for example, under the assumption that CTCF molecules bound at BEs have an intrinsic affinity for each other. Note that, in contrast with experimental data, strong peaks are observed between all possible pairs of BEs in this model, and not preferentially between BEs at short separations along the linear genome. The outcome of this simulation suggests something more than long-range looping-mediated interaction is required to form TADs and peaks; in particular, an additional mechanism is required to restrict interactions to occur between proximal elements along the linear genome. **c.** *top*: schematic of the model, chromatin fiber in grey, BEs in orange. *bottom*: arc-diagram showing BE-to-BE interactions realized in this conformation (solid lines), and all possible BE-to-BE interactions (dashed lines). **d.** contact map and *P(s)* for direct BE-to-BE model with attraction strength 1.5kT. Note the peaks in contact probability between non-proximal boundary elements (pink circles) as well as between proximal elements (black circles). **e.** attraction strength 3kT. **f.** attraction strength 5kT.

**Fig. S12.**
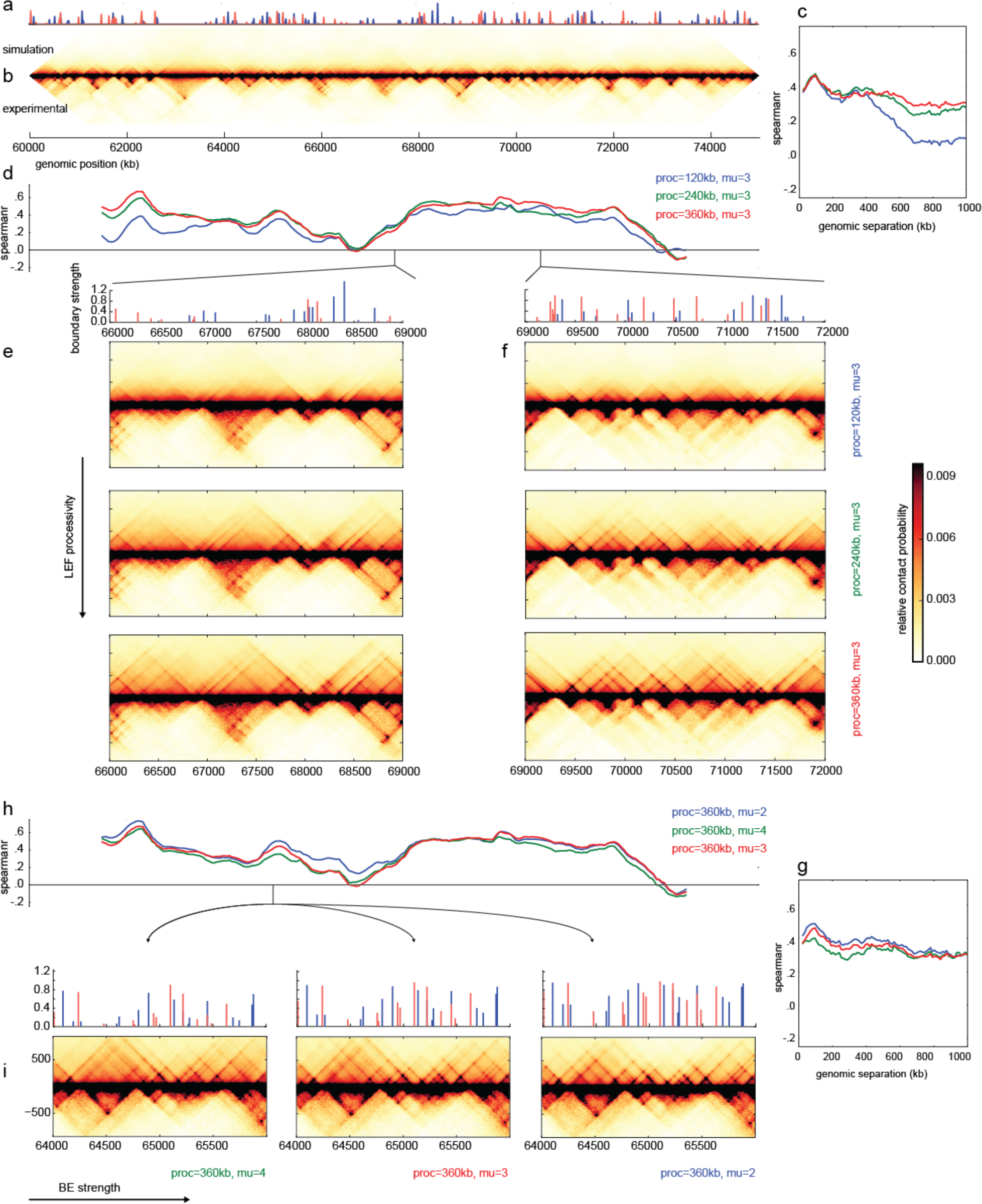
Model with orientation-specific BEs of varying permeability. **a.** Orientation-specific BE strength profile across the simulated region of human chromosome 14, cell type GM12878, 12kb resolution. BE strength reflects the average number of BEs bound within a 12kb (20 monomer) region. **b.** Simulated contact map of the full region of human chr14, GM12878 cell type. Maps are compared with experimental maps for the same regions at the same 12kb resolution. **c.** Spearman correlation between experimental and simulated observed/expected contact maps as a function of genomic distance for simulations with BE parameter mu=3, and LEF processivity 120kb (blue), 240kb (green), and 360kb (red). Note this corresponds to correlation between the maps at increasing distances from the center line of the maps. **d.** Correlation profile between experimental and simulated observed/expected along the chromosome for sliding genomic windows of 3Mb along the chromosome by 1Mb maximum separation. **e. f.** Simulated contact maps for two indicated regions with BE parameter mu=3, and LEF processivity 120kb, 240kb, and 360kb. **g.** Spearman correlation between experimental and simulated observed/expected contact maps as a function of genomic distance for simulations with LEF processivity 360kb for BE parameter mu= 2 (blue), 4 (green), 3 (red). **h.** Correlation profile between experimental and simulated observed/expected (as in d). **i.** Simulated contact maps for two indicated regions with LEF processivity 360kb for BE parameter mu= 2, 3, 4.

**Fig. S13.**
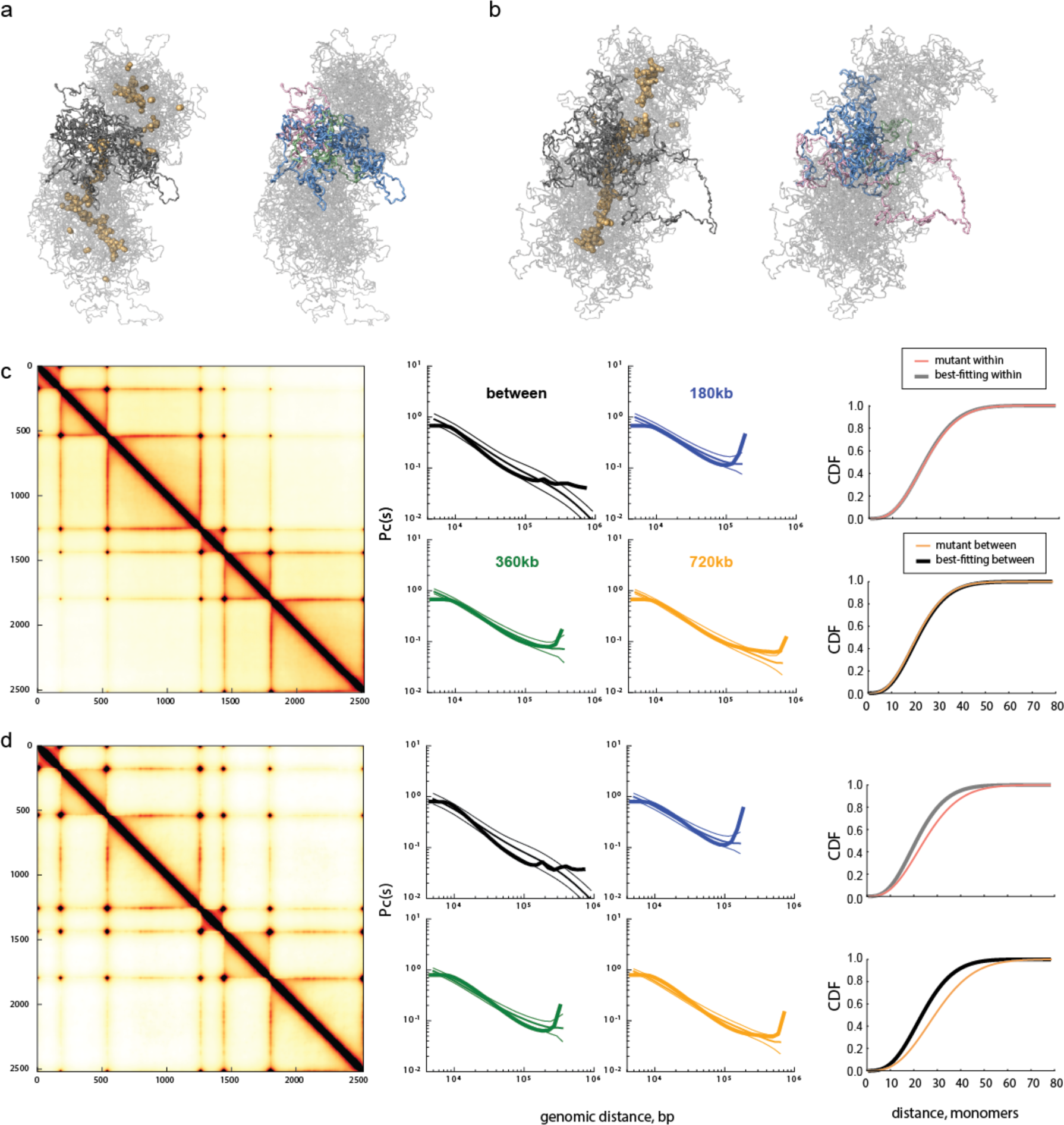
Increased LEF lifetime creates a prophase-like vermicelli state. **a.** Conformations of a polymer subject to LEF dynamics for very high LEF processivity (1200kb), ten-times the processivity in **Fig 1E**; such high processivity might occur following depletion of LEF-unloading factors. *Left*: shows LEFs (yellow spheres), and chromatin (grey), *right*: shows three neighboring regions between BEs of sizes (180kb, 360kb, 720kb) in (green, pink, blue) for the same conformation (as colored in **Fig 1D**). Interestingly, LEFs form a continuous backbone, despite the presence of boundaries. **b.** As previously, but for even larger processivity (6000kb). **c.** Processivity, separation = 1200kb, 120kb (as for **a**), *Left*: contact map, *Middle*: associated *P(s)*for three TAD sizes and between TADs. *Right*: CDF. 3D-to-1D ratio 5000/4, stiffness 2, density .2, cutoff 8. **d.** as for c, but with larger processivity (6000kb). Interestingly, the results in (c,d) contrast with the result (**fig. S7**) that increasing the processivity to 480kb (4-fold) from the best-fitting model decreases FISH distances. Indeed, despite large changes on the contact map, the processivity 1200kb (10-fold increased) simulated FISH CDF are almost unchanged from the best-fitting model with 120kb, and the processivity 6000kb CDF actually shifts to larger distances. This behavior arises because the simulated FISH CDF measured at 360kb reflects both the overall compaction, as well as the average loop size. While the overall compaction of the chromosome is certainly higher for 1200kb and 6000kb than for 480kb, the average loop size is also higher due to the increased formation of nested loops (Goloborodko et al., 2015), increasing the loop size in this very dense regime increases the average distance of non-BE monomers to the LEF backbone, which in turn increases distances between any two non-BE monomers. We note this is analogous to how over-condensed mitotic chromosomes are thicker and shorter than normal mitotic chromosomes. Together, these results indicate the importance of measuring compaction at multiple scales for testing predictions of the loop extrusion mechanism.

**Fig. S14.**
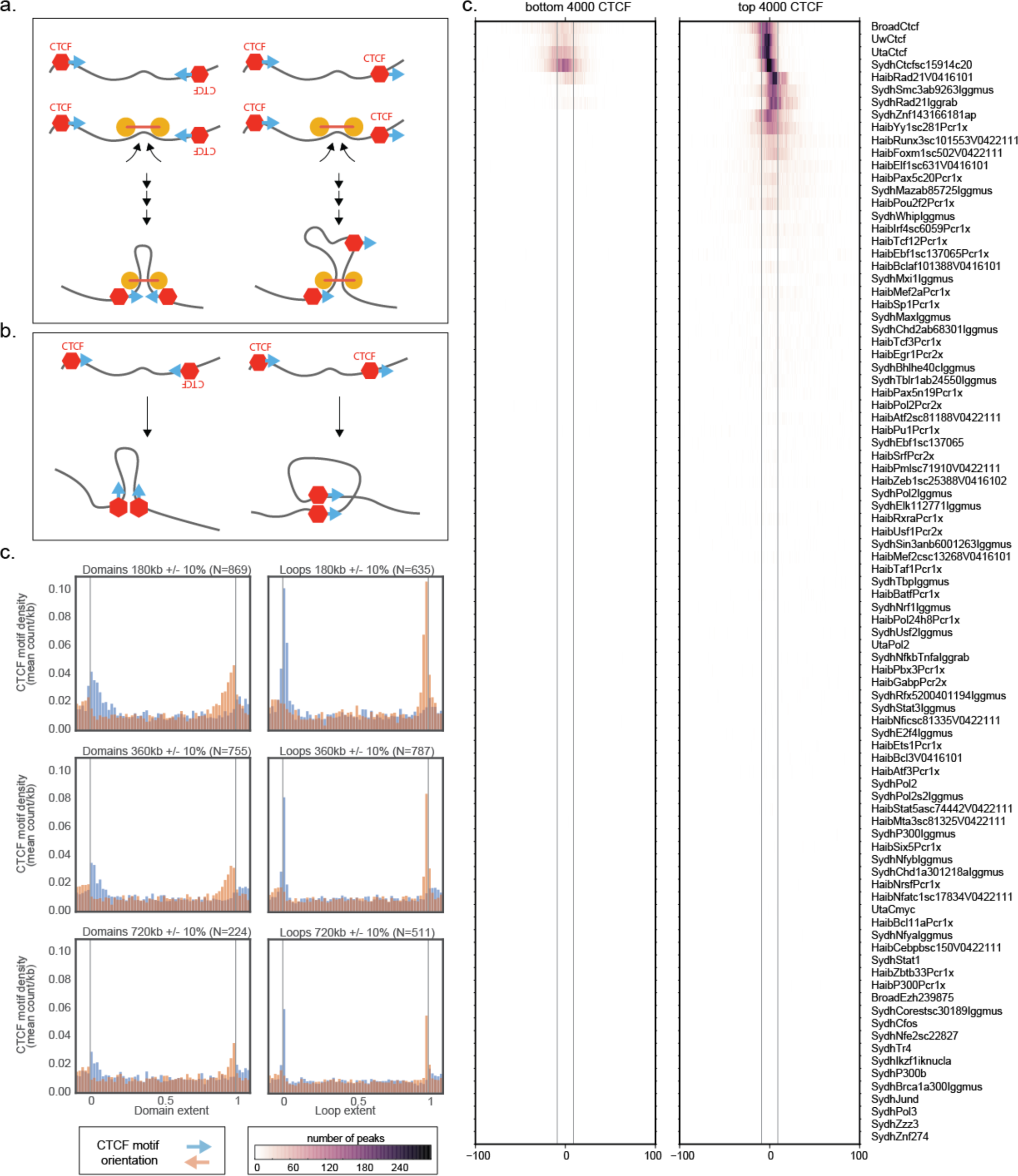
Loop extrusion can maintain bound CTCF orientation. Genomics experiments (Rao et al., 2014; Rudan et al., 2015) have revealed preferential enrichment of inwardly oriented CTCF motifs at TAD boundaries and between peak locus pairs; this has been taken to suggest that CTCF can stabilize chromatin loops in an orientation-dependent manner. We find that preferential enrichment persists over a range of TAD sizes and peak locus pair genomic distances (see below). Here we illustrate how two mechanisms of loop formation would correspond to motif orientations. The first mechanism relies on loop extrusion, and in the second, two looping elements pair after a stochastic encounter in 3D. We illustrate how loop extrusion can lead to the observed orientation-specific enrichment, while 3D encounters cannot. **a.** loop extrusion maintains the orientation of motifs over large separation, whether for convergent motif-pairs (*left*), or for same-orientation motif-pairs (*right*). If CTCFs halt loop LEF extrusion and stabilize loops in an orientation-dependent manner, then the mechanism of loop extrusion explored here can explain the observed enrichment in convergent CTCF motifs at TAD boundaries and loop bases, even at very large genomic separations. **b.** encounters in 3D at genomic distances much larger than the twist persistence length of the chromatin fiber would maintain no preferential orientation of motif-pairs. **c.** (*left*) enrichment plot of CTCF motif orientation (similar to (Rudan et al., 2015)) over TADs called in GM12878 (Rao et al., 2014), for TADs of three sizes: 180kb, 360kb, and 720kb, with a +/-10% size range. Each TAD in a given size range was divided into 50 bins, and then extended by 5 bins to the left and right. The total number of + or - motif occurrences in each bin was aggregated, and divided by the bin size to give a motif ״density״ per kb. The bar heights give the average motif density in each bin. Motif occurrences and orientations were obtained from a publicly-available resource of regulatory motifs derived from a combination of known literature motifs and motifs discovered from ENCODE TF ChIP-seq data (Kheradpour and Kellis, 2014). We selected motif matches corresponding to factor group “CTCF_known1” passing a position weight-matrix based p-value threshold. (*right*) similarly, but for peak loci (also called loops, in (Rao et al., 2014)).Note that motif orientation enrichment is similar over each range of distances, with TADs displaying a ∼2-fold and loops ∼3- to 5-fold enrichment of inward-facing motifs at each range of distances. **d.** histograms of GM12878 ChIP-seq peak summits around the 4000 weakest (left) and 4000 strongest (right) CTCF core motif instances (CTCF_known1, consensus 5‘-TGGCCACCAGGGGGCGCTA-3’) for all ENCODE transcription factors (uniformly processed narrowPeak files from all production centers) (ENCODE consortium, 2012). The strength of a motif instance was determined by the maximal fold change signal of the CTCF ChIP-seq peak overlapping the motif. The histograms span a 200-bp window and are aligned along the center of the core motif, whose width is depicted with vertical grey lines. The histograms are oriented along the reference stand of the corresponding motif instance. The strongest enrichment is seen for CTCF, SMC3, RAD21, ZNF143 and YY1. Note that the cohesin subunit (SMC3 and RAD21) densities are shifted downstream relative to those of the other factors.

## Movie and Database D1 Captions

### Movie-M1

This movie illustrates dynamics of a 120-kb TAD in a best-fitting model (**Fig 1E**). 320 consecutive simulation blocks (i.e. rounds of 3D-simulation time-steps followed by 1D-simulation time-steps) are shown, with 6 linearly interpolated conformations between any two blocks. Loop bases are shown as spheres, and regions within loops are shown in black.

### Movie-M2

Modification of the **Movie M1** for the model with twice-longer processivity, 240 kb compared to 120kb in the **Movie M1.** Monomers in loops are highlighted by black. For this value of processivity, loops-inside-loops occurred frequently. Monomers in the inside of multiple loops are highlighted by purple. Note that when the outer loop in a loop-in-loop conformation disappears, purple changes to black.

### Database D1

The database contains goodness-of-fit for all 6192 parameter-sets.

